# Modeling reveals human-rodent differences in h-current kinetics influencing resonance in cortical layer 5 neurons

**DOI:** 10.1101/2020.02.12.945980

**Authors:** Scott Rich, Homeira Moradi Chameh, Vladislav Sekulic, Taufik A. Valiante, Frances K. Skinner

**Affiliations:** Division of Clinical and Computational Neuroscience, Krembil Research Institute, University Health Network (UHN), Toronto, Ontario, Canada; Institute of Biomaterials and Biomedical Engineering, University of Toronto, Toronto, ON, Canada; Institute of Medical Science, University of Toronto, Toronto, ON, Canada; Division of Neurosurgery, Department of Surgery, University of Toronto, Toronto, ON, Canada; Electrical and Computer Engineering, University of Toronto, Toronto, ON, Canada; Departments of Medicine (Neurology) and Physiology, University of Toronto, Toronto, ON, Canada

**Keywords:** Computational model, h-current, cortical layer 5, subthreshold resonance, human neurons

## Abstract

While our understanding of human neurons is often inferred from rodent data, inter-species differences between neurons can be captured by building cellular models specifically from human data. This includes understanding differences at the level of ion channels and their implications for human brain function. Thus, we here present a full spiking, biophysically-detailed multi-compartment model of a human layer 5 (L5) cortical pyramidal cell. Model development was primarily based on morphological and electrophysiological data from the same human L5 neuron, avoiding confounds of experimental variability. Focus was placed on describing the behavior of the hyperpolarization-activated cation (h-) channel, given increasing interest in this channel due to its role in pacemaking and differentiating cell types. We ensured that the model exhibited post-inhibitory rebound (PIR) spiking considering its relationship with the h-current, along with other general spiking characteristics. The model was validated against data not used in its development, which highlighted distinctly slower kinetics of the human h-current relative to the rodent setting. We linked the lack of subthreshold resonance observed in human L5 neurons to these human-specific h-current kinetics. This work shows that it is possible and necessary to build human-specific biophysical neuron models in order to understand human brain dynamics.

## Introduction

Currently, much of what is understood about specific cell-types and their role in “computation” (Womelsdorf et al., 2014) within the six-layered neocortex stems from invasive and *in vitro* studies in rodents and non-human primates. Whether or not such principles can be extended to human neocortex remains speculative at best. Despite the significant transcriptomic convergence of human and mouse neurons (Hodge et al., 2019), significant differences between human and rodent cell-type properties exist. *In vitro* studies have identified differences between mouse and human neurons in morphology (Mohan et al., 2015), dendritic integration (Beaulieu-Laroche et al., 2018; Eyal et al., 2016), synaptic properties (Verhoog et al., 2013), and collective dynamics (McGinn and Valiante, 2014; Molnár et al., 2008; Florez et al., 2013). However, less explored are the active, voltage-gated ion channels of human cortical neurons. The hyperpolarization-activated cation (h-) channel is a voltage-gated channel of increasing interest, considering recent findings showing that its expression may serve to differentiate cell types (Nandi et al., 2020). Additionally, it has long been attributed with a pacemaking role and hence of potential importance in functionally relevant brain rhythms (Akam and Kullmann, 2014; Buzsaki, 2006; Womelsdorf et al., 2014; Fries, 2005; Anastassiou et al., 2011; Hanslmayr et al., 2019; Vaz et al., 2019).

The presence of h-channels in a neuron leads to a characteristic “sag” in voltage recordings when hyperpolarizing current steps are applied. A larger sag implies a larger current through the h-channel (h-current), and is often attributed to a larger channel conductance. Recently it has been shown that increased expression of h-channels contribute to the observed subthreshold resonance in supragranular layer human pyramidal cells not seen in their rodent counterparts (Kalmbach et al., 2018). Such differential expression of h-channels also appears to be present between superficial and deep layer neurons of human cortex, with layer 5 (L5) pyramidal cells demonstrating a larger sag mediated by h-channels when compared to those in layer 2/3 (L2/3) (Chameh et al., 2019). However, despite the presence of large sag currents in human L5 pyramidal cells, they very rarely exhibit subthreshold resonance (Chameh et al., 2019). This result is surprising considering recent human work in L2/3 (Kalmbach et al., 2018), as well as findings that rodent L5 pyramidal cells commonly exhibit subthreshold resonance (Silva et al., 1991; Ulrich, 2002; Dembrow et al., 2010; Schmidt et al., 2016).

The implied relationship between the h-current and subthreshold resonance is of great interest: not only is this dynamic well-studied experimentally (Kalmbach et al., 2018; Hu et al., 2002, 2009; Zemankovics et al., 2010; Ulrich, 2002), but the emergence of brain oscillations has a potential, albeit unclear, dependence on subthreshold resonance (Richardson et al., 2003; Rotstein, 2017). Relating subthreshold resonance to spiking and brain oscillations is a complicated endeavour that necessarily depends on the entirety of a neuron’s characteristics, including its full complement of voltage-gated channels, passive properties, and morphology. Concerted efforts to understand these relationships are the subject of ongoing theoretical research (Rotstein and Nadim, 2014; Rotstein, 2014). Toward this end, untangling the relationship between the h-current and subthreshold resonance is crucial, especially considering the unexpected results reported by Chameh et al. (2019). Unfortunately, while the experimental literature (Kalmbach et al., 2018; Chameh et al., 2019) has revealed that differences in the capacity for subthreshold resonance exist between human and rodent neurons, the reason(s) *why* these differences arise remains unclear. Computational modelling is required to address such questions.

While such computational investigations would be particularly suited to dissect these unanticipated results, at present we lack human neuron models with the same level of biophysical detail as rodent models. The development of detailed models of human neurons, including precise articulations of the dynamics of voltage-gated channels and experimentally-constrained morphologies, are more challenging than their rodent counterparts due to limited access to tissue for experimental recordings. For example, one cannot control for age or gender in acquiring human tissue. This challenge is exacerbated by the fact that one should obtain a complete experimental data set of morphology and electrophysiological recordings from the same neuron to best constrain a model. Doing so avoids obtaining an incorrect “balance” of channel contributions due to averaging across data sets given biological variability (Marder and Goaillard, 2006; Marder and Taylor, 2011) considering that conductances in similarly classified cells can vary 2-6 fold (Goaillard et al., 2009; Ransdell et al., 2013).

Easier access to rodent tissue makes obtaining this type of data more feasible, and massive amounts of rodent data are becoming available (Dong, 2008; Jones et al., 2009; Sunkin et al., 2012). Such data, together with the development of model databases, can uncover degeneracies and expose compensatory biophysical mechanisms for neuron and network dynamics (Marder and Taylor, 2011; Prinz et al., 2004). However, this relies on the existence of large sets of data and computational models, both of which are not yet present for human neurons. Despite these limitations, the clear differences between human and rodent neurons (Hodge et al., 2019; Eyal et al., 2018, 2016; Testa-Silva et al., 2014; Verhoog et al., 2013; Beaulieu-Laroche et al., 2018) necessitates that we do create human neuron models to help us understand human brain dynamics. The challenge, then, is how to make best use of the available data from human neurons to create models in this particular setting.

In this work we take advantage of a novel set of data that includes a morphological reconstruction, current clamp recordings, and voltage clamp recordings, obtained from the *same* human L5 pyramidal cell. In the absence of particular information in the human setting, we include in our model the same ion channel types and distributions used in a similarly detailed rodent L5 pyramidal cell model (Hay et al., 2011). Using this data, we design a modeling strategy that primarily constrains the neuron’s passive and active properties based on current clamp data. Considering the data contains numerous current clamp recordings at hyperpolarized voltages, we exploit the fact that h-channels are the primary active voltage-gated channels at hyperpolarized voltages (Toledo-Rodriguez et al., 2004) to model its activity in great detail on the backbone of the reconstructed neuron. Simultaneously, we restrict the relative contributions of the additional ion channel types influencing the neuron’s dynamics at voltages near or above the resting membrane potential, along with its passive properties. As these recordings were obtained in the presence of a voltage-gated sodium channel blocker, spiking was not present in this data set. Correspondingly, we analyzed post-inhibitory rebound (PIR) spiking and repetitive firing characteristics from another set of human L5 neurons (Chameh et al., 2019) to impart constraints on the model’s spiking characteristics. A particular emphasis is placed on ensuring our model neuron exhibits reasonable PIR spiking behavior, a dynamic which is shown by Chameh et al. (2019) to be reliant upon h-channels in human L5 neurons.

This process yields a full spiking, biophysically detailed, multi-compartment model of a distinctly human L5 cortical pyramidal cell. The resulting model matches the electrophysiological data from hyperpolarizing current clamp experiments in the primary cell remarkably well, while also demonstrating repetitive PIR spiking properties characteristic of human L5 pyramidal cells from Chameh et al. (2019). Further, our model makes possible a comparison of these neurons in the human and rodent setting, revealing important differences in both the conductance and kinetics of the h-current across species.

The decision to more precisely model the activity of the h-current is motivated by the clear preponderance of h-channels in human L5 neurons (Chameh et al., 2019), their complex role in regulating neuronal excitability (Dyhrfjeld-Johnsen et al., 2009; Biel et al., 2009), their role in dictating cell types (Nandi et al., 2020), and their hypothesized role in driving subthreshold resonance (Kalmbach et al., 2018; Hu et al., 2002, 2009; Zemankovics et al., 2010; Ulrich, 2002). Our neuron model contains a new articulation of the activity of the h-current in the human setting, derived entirely by “fitting” the model to current clamp data. This “human” h-current model is validated against experimentally-derived kinetics from voltage clamp data not used as a modeling constraint. Pivotally, these kinetics are distinct from those observed in rodents and implemented in many rodent cortical pyramidal cell models (Kole et al., 2006). Subthreshold resonance characteristics provided an additional avenue by which to validate our model to experimental data, and indeed our model lacks subthreshold resonance just as observed experimentally (Chameh et al., 2019). This additional validation justifies a detailed investigation into the generation of subthreshold resonance in these neurons, which reveals that the unique kinetics of human h-currents explain the lack of resonance seen in human L5 pyramidal cells despite the abundance of these channels indicated by large sag currents (Chameh et al., 2019).

In summary, our model development strategy and the resulting human neuron model answer precise questions about the role of h-channels in the different dynamics exhibited by similarly classified human and rodent neurons. Specifically, our findings reveal that the important differences in dynamics of h-currents in human L5 pyramidal cells, when compared to their rodent counterparts, obviate subthreshold resonance at resting membrane potential despite the presence of large sag currents. Given the numerous ways in which the validity of the model is confirmed, along with the implementation of spiking behaviors, our model has the potential to be used in future studies to disentangle the intricacies underlying other functionally important dynamics, including suprathreshold behaviors such as the frequency-dependent gain (Higgs and Spain, 2009) of these neurons. The multitude of ways in which this specific model can be utilized indicates that the modeling strategy presented here may be generally applicable. This publicly available cell model represents the first biophysically detailed, multi-compartment human L5 pyramidal cell model with a full complement of ion channel types and distributions designed to investigate distinctly human neural dynamics.

## Methods and Materials

### Experimental recordings of human L5 cortical pyramidal cells

We note that the experimental protocol described below is *distinct* from that in Chameh et al. (2019), in which (as examples) the data was collected without synaptic blockers and without tetrodotoxin (TTX). The protocol used in this study was designed specifically with the goal of creating a detailed neuron model in mind. This motivated the desire to get the maximal amount of data possible from the same neuron to avoid biological variability and inappropriate characterization from averaging (Marder and Taylor, 2011). The protocol was also designed to facilitate our characterization of the h-current with the morphology and passive properties of this particular neuron. While this choice is well-rationalized, there are limits to the amount of applicable data that can be obtained from the same neuron. These limitations informed the modeling strategy described below. Access to human tissue provided no control over age, gender, or the particular aspect of the surgery involved, which only adds to the issue of experimental variability in recording between similarly classified cells and cements the importance of prioritizing data obtained from a single neuron in model generation.

#### Ethics statement

Surgical specimens were obtained from Toronto Western Hospital. Written informed consent was obtained from all study participants as stated in the research protocol. In accordance with the Declaration of Helsinki, approval for this study was received by the University Health Network Research Ethics board.

#### Acute slice preparation from human cortex

Neocortical slices were obtained from the middle temporal gyrus in patients undergoing a standard anterior temporal lobectomy for medically-intractable epilepsy (Mansouri et al., 2012). Tissue obtained from surgery was distal to the epileptogenic zone tissue and was thus considered largely unaffected by the neuropathology. We note that this is the same area from which recent data characterizing human L3 cortex was obtained (Kalmbach et al., 2018).

Immediately following surgical resection, the cortical block was placed in an ice-cold (approximately 4°C) slicing solution containing (in mM): sucrose 248, KCl 2, MgSO4.7H2O 3, CaCl2.2H2O 1, NaHCO3 26, NaH2PO4.H2O 1.25, and D-glucose 10. The solution was continuously aerated with 95% O2-5% CO2 and its total osmolarity was 295-305 mOsm. Tissue blocks were transported to the laboratory within 5 min. Transverse brain slices (400 *μ*m) were cut using a vibratome (Leica 1200 V) in slicing solution. Tissue slicing was performed perpendicular to the pial surface to ensure that pyramidal cell dendrites were minimally truncated (Kostopoulos et al., 1989; Kalmbach et al., 2018). The slicing solution was the same as used for transport of tissue from the operation room to the laboratory. The total duration of transportation and slicing was kept to a maximum of 20 minutes, as suggested by Köhling and Avoli (2006).

After sectioning, the slices were incubated for 30 min at 34°C in standard artificial cerebrospinal fluid (aCSF). The aCSF contained (in mM): NaCl 123, KCl 4, CaCl2.2H2O 1, MgSO4.7H2O 1, NaHCO3 25, NaH2PO4.H2O 1.2, and D-glucose 10, pH 7.40. All aCSF and slicing solutions were continuously bubbled with carbogen gas (95% O2-5% CO2) and had an osmolarity of 295-305 mOsm. Following this incubation, the slices were kept in standard aCSF at 22–23°C for at least 1 h, until they were individually transferred to a submerged recording chamber.

#### Electrophysiological recordings

##### Experimental setting

*In vitro* whole-cell recordings were obtained from human neocortical L5 neurons. For recording, slices were transferred to a recording chamber mounted on a fixed-stage upright microscope (Axioskop 2 FS MOT; Carl Zeiss, Germany), and were continually perfused at 8 ml/min with standard aCSF at 32-34 oC. All experiments were performed with excitatory (APV 50 *μ*M, Sigma; CNQX 25 *μ*M, Sigma) and inhibitory (Bicuculline 10 μM, Sigma; CGP-35348 10 *μ*M, Sigma) synaptic activity blocked. Cortical neurons were visualized using an IR-CCD camera (IR-1000, MTI, USA) with a 40x water immersion objective lens. Upon visual inspection, L2/3 of the cortex is differentiated from L5 by a tight band of densely packed cells representing layer 4 (L4). L5 of the cortex is much less dense than L4, and contains notably larger pyramidal cells, providing us with the necessary confidence that these experiments were performed on L5 neurons.

Patch pipettes (3-6 MΩ resistance) were pulled from standard borosilicate glass pipettes (thin-wall borosilicate tubes with filaments, World Precision Instruments, Sarasota, FL, USA) using a vertical puller (PC-10, Narishige). Pipettes were filled with intracellular solution containing (in mM): K-gluconate 135, NaCl 10, HEPES 10, MgCl2 1, Na2ATP 2, GTP 0.3, and biocytin (3-5mg/mL). The solution’s pH was adjusted with KOH to 7.4 and its osmolarity was 290–300 mOsm. Whole-cell patch-clamp recordings were obtained with an Multiclamp 700A amplifier and pClamp 9.2 data acquisition software (Axon instruments, Molecular Devices, USA). Subsequently, electrical signals were digitized at 20 kHz using a 1320X digitizer. The access resistance was monitored throughout the recording (typically between 8-25 MΩ), and cells were discarded if access resistance was > 25 MΩ. The liquid junction potential was calculated to be 10.8 mV which is corrected for whenever the experimental data is used for modeling or in direct comparison to model values (i.e. Figure 6), but not when the experimental data is presented on its own (i.e. Figures 7**B** and 10**E-F**).

##### Current clamp data

Current clamp data from the primary neuron is used as the primary constraint for the computational model. This data was obtained in the following fashion. Hyperpolarizing current pulses (1000 ms duration, −50-400 pA, step size: 50 pA) and depolarizing current pulses (1000 ms duration, 50-400 pA/ step size: 50 pA) were injected to measure passive and active membrane properties in presence of voltage gated sodium channels blocker (TTX 1 *μ*M; Alomone Labs). This data is highlighted in Figure 1.

**Figure 1.**
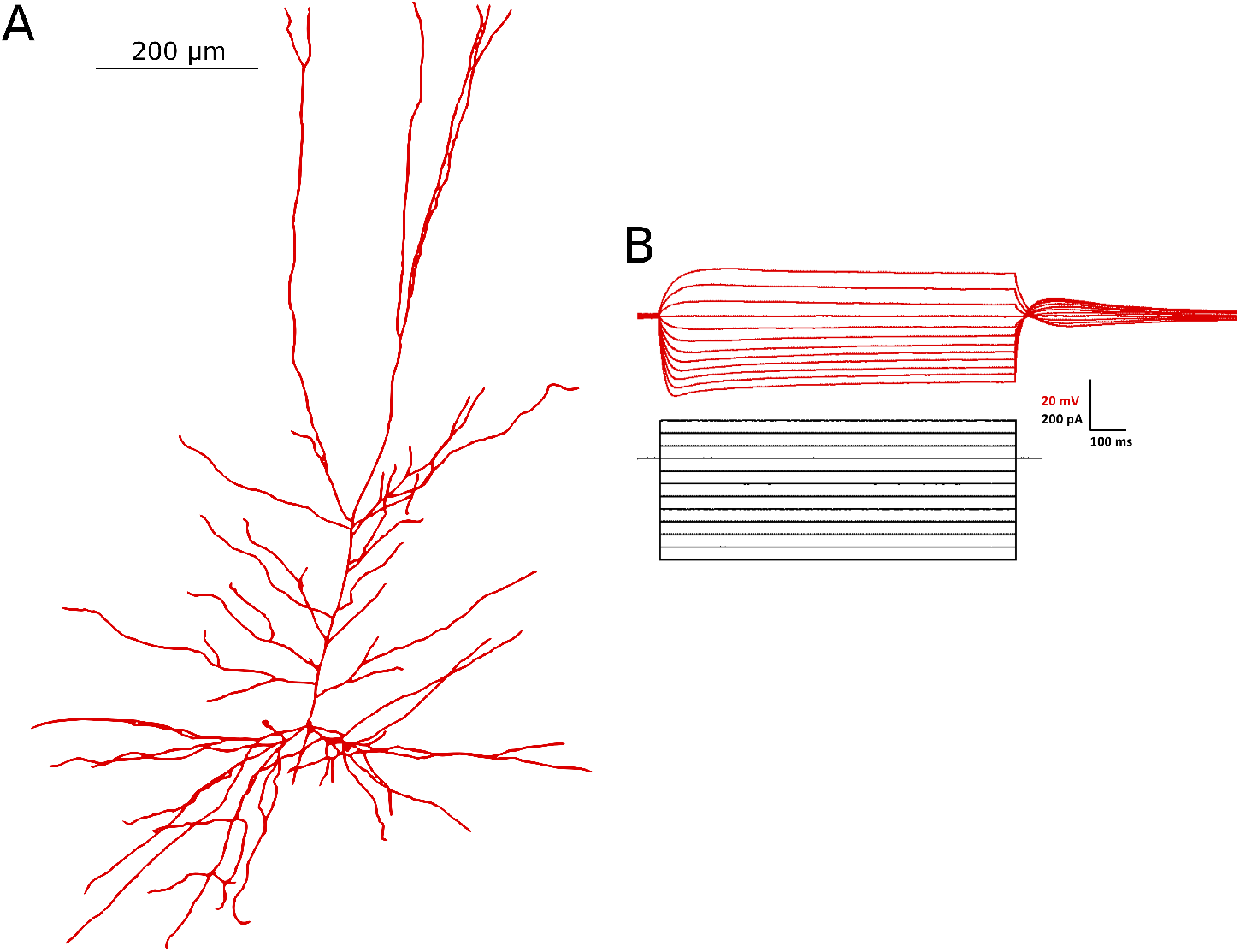
Morphology and current clamp data obtained from the primary neuron. **(A)** The morphology of the primary neuron was reconstructed using IMARIS software and imported into NEURON (which generated the plot shown here). For visualization purposes, a uniform diameter is shown for each compartment in this image. **(B)** Current clamp recordings from the primary neuron in the presence of TTX that are the primary constraining data for model development.

##### Voltage clamp data

To characterize the h-current kinetics, 1000 ms-long voltage clamp steps were used in −10 mV increments, down to −140 mV from a holding potential of −60 mV. The tail current was quantified as the difference between peak amplitude of residual current at the end of each holding potential and the steady state current from holding potentials of −140 to −60 mV. This value was used to calculate the steady-state activation curve as presented in Figure 6 by normalizing these values between 0 and 1. To calculate the h-current time constant, a single or double-exponential model was fitted to the initial response of the neuron to the voltage clamp using Clampfit 10.7 (Axon Instruments, Molecular Devices, USA). In experiments quantifying the h-current kinetics, tetrodotoxin (TTX, 1 μM; Alomone Labs) to block voltage gated sodium currents, CoCl2 (2mM; Sigma-Aldrich) to block voltage-sensitive calcium currents, and BaCl2 (1mM; Sigma-Aldrich) to block inwardly rectifying potassium current were added to the bath solution. These recordings were taken both in the primary cell and in a secondary set of L5 pyramidal cells, the data for both of which are presented in Figure 6.

##### Spiking data obtained from a separate neuron population

Spiking data from L5 neurons are obtained from part of the population of neurons characterized by Chameh et al. (2019). The data used in this work, as presented in Figure 4**E** and **F**, was obtained from 76 cells. In Figure 4**F** only cells that exhibited a PIR spike are shown.

##### Modeling motivations drive experimental protocol design

We focused our efforts on obtaining a single primary neuron from which we could obtain both a reliable morphology and a large suite of electrophysiological data (in both current clamp and voltage clamp modes). Several attempts were required to successfully accomplish this task. We note that TTX was applied to our primary neuron in order to ensure that the potential of the cell to spike under mild current clamp perturbations did not interfere with our ability to accurately capture the neuron’s passive properties; while passive properties are ideally calculated under a full blockade of ion channels, the use of just TTX allowed us to balance the goals of accurately encapsulating both the neuron’s passive properties and the influence of the h-current and other ionic currents implemented in the model.

We further note that given time limitations to our experimental protocol imposed by the use of human tissue, we were unable to perform voltage clamp experiments both with and without the h-channel blocker ZD in the same cell to truly “isolate” the h-current. We thus decided to use current clamp data to constrain our model; along with space-clamp issues, without ZD recordings we cannot assert with full certainty that the h-current features derived from voltage clamp data are not influenced by other channels. It is for this reason that this data was instead used for model validation, in which these “approximate” values of the h-current kinetics are more appropriate, rather than direct model constraint.

#### Histological methods and morphological reconstruction

During electrophysiological recording, biocytin (3-5 mg/ml) was allowed to diffuse into the patched neuron; after 20-45 min, the electrodes were slowly retracted under visual guidance to maintain the quality of the seal and staining. The slices were left for another 10-15 min in the recording chamber to washout excess biocytin from the extracellular space, then slices containing biocytin-filled cells were fixed in 4% paraformaldehyde in PBS for 24 hours at 4°C. The slices were washed at least 4×10 min in PBS solution (0.1 mM). To reveal biocytin, slices were incubated in blocking serum (0.5% Bovine serum albumin (BSA), 0.5% milk powder) and 0.1% Triton X-100 in PBS for 1 hour at room temperature.

Finally, slices were incubated with streptavidin fluorescein (FITC) conjugated (1:400) (Thermo Fisher Scientific, Canada) on a shaker at 4°C for 12 hours. Then slices were washed at least 4×10 min in PBS and mounted on the glass slide using moviol (Sigma-Aldrich). Imaging was done using a Zeiss LSM710 Multiphone microscope. Cellular morphology was reconstructed using IMARIS software (Bitplane, Oxford instrument company). These steps were performed on the same neuron from which the current clamp traces were obtained, yielding the morphology shown in Figure 1. The number of compartments in the final reconstruction of the primary human L5 pyramidal cell was 211. This was verified to be numerically appropriate in simulations performed.

#### Subthreshold resonance

Experimental data regarding the subthreshold resonance of L5 neurons was first presented in Chameh et al. (2019), with an example trace replicated here in Figure 7**B**. We include the method by which this data was obtained here for completeness.

To assess the subthreshold resonance properties of L5 pyramidal cells, a frequency modulated sine wave current input (ZAP) was generated ranging from 1 to 20 Hz, lasting 20 s (Hutcheon et al., 1996) with a sampling rate of 10 kHz. This current waveform was then injected using the custom waveform feature of Clampex 9.2 (Axon Instruments, Molecular devices, USA) at either the neuron’s resting membrane potential or around a held hyperpolarized voltage. The subthreshold current amplitude was adjusted to the maximal current that did not elicit spiking. The impedance curve resulting from this experiment was calculated as illustrated by Puil et al. (1986). Summarized briefly, the impedance is calculated by dividing the power spectrum of the voltage trace by the power spectrum of the current trace under a ZAP protocol. Given the noisiness of these plots, in our presentations we also include a “smoothed” version of these curves simply calculated using the *smooth* function in MATLAB (MATLAB, 2019).

### Construction of multi-compartment computational model of a human L5 cortical pyramidal cell

The code containing the final model, as well as various tools to perform *in silico* experiments, can be found at https://github.com/FKSkinnerLab/HumanL5NeuronModel. We describe the development of this model below.

#### Morphological reconstruction and ionic currents

Our model generation process began with a reconstruction of the primary neuron’s cellular morphology, illustrated in Figure 1, and implementation of this reconstruction in the NEURON simulation environment (Carnevale and Hines, 2006). In the absence of specific knowledge of the various ion channel types and their distributions in the human setting, we included ten different types of ion channels that were used in developing rodent L5 pyramidal cell models (Hay et al., 2011). Thus the human L5 pyramidal cell model, before any adjustments or parameter optimization, consisted of the same 10 ion channel types producing the ionic currents present in the multi-compartment model.

The ion channels include the following: a fast, inactivating sodium current (abbreviated Na_Ta); a persistent sodium current (abbreviated Nap_Et2); a slow, inactivating potassium current (abbreviated K_Pst); a fast, non-inactivating potassium current (abbreviated SKv3_1); a small-conductance calcium activated potassium current (abbreviated SK_E2); a fast, inactivating potassium current (abbreviated K_Tst); a low-voltage activated calcium current (abbreviated Ca_LVA); a high-voltage activated calcium current (abbreviated Ca_HVA); the non-specific hyperpolarization-activated cation current which we refer to as the h-current (abbreviated Ih); and the voltage-gated muscarinic potassium channel (abbreviated Im). Note that the abbreviations used here are motivated by the labeling used in the NEURON code for consistency.

#### Mathematical equations and parameter values

The mathematical equations describing the currents used a conductance-based formalism as given in the Methods of Hay et al. (2011). They were used unaltered except for Ih, Na_Ta, and SKv3_1.

The Ih kinetics were fit from scratch to allow for any potential differences between rodent and human h-currents to be captured. We used a general mathematical model structure as used in previous work to model h-current dynamics (Sekulić et al., 2019) and included the parameters in this model in our optimizations.

The equations for the h-current model are as follows:

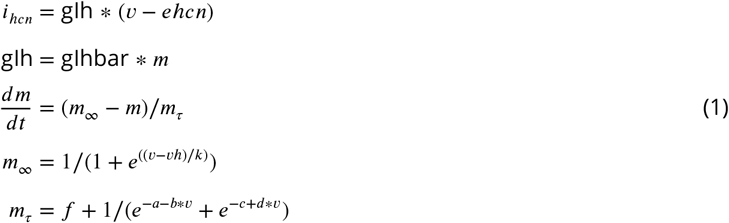

where *i_hcn_* is the current flow through the h-channels (mA/cm^2^), gIh is the conductance (S/cm^2^), *v* is the voltage (mV), gIhbar is the maximum conductance (S/cm^2^) (an optimized parameter), *m* is the unitless gating variable, *t* is time (ms), *vh* is the half-activation potential (mV) (an optimized parameter), *ehcn* is the reversal potential (mV) (an optimized parameter) *k* is the slope of activation (an optimized parameter), and *a, b, c, d* and *f* are optimized parameters (ms). *m*_oo_ is the steady-state activation function and *m_r_* is the time constant of activation.

The changes to the Na_Ta and SKv3_1 ionic currents were simple “shifts” of the activation curves to more hyperpolarized voltages, as necessitated to best replicate experimentally measured PIR characteristics of human L5 cortical pyramidal cells as in Figure 4**F**.

The specific equations where these changes are implemented are shown below:

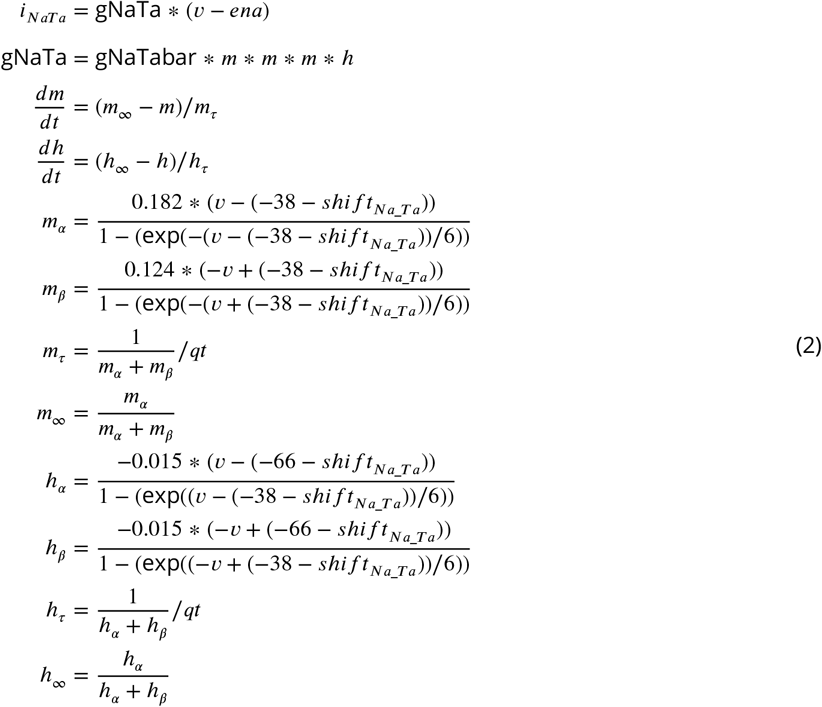

where *qt* is a local constant equal to 2.3^(34−21)/10^ ;

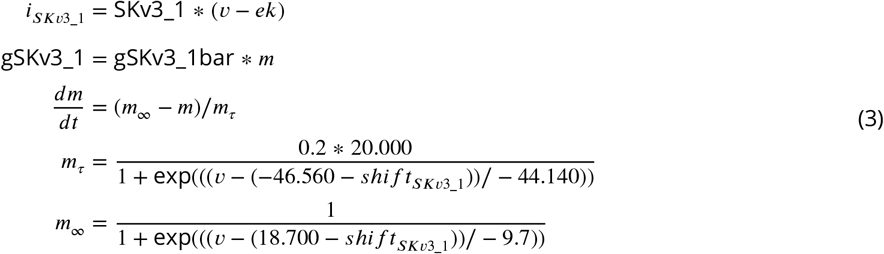

The units of the *i* (current), *g* (conductance), *v* (voltage), *e* (reversal potential), and *t* (time) terms in both of these equations are as given above for the h-current. *ena* refers to the reversal potential of sodium and *ek* refers to the reversal potential of potassium, both of which are unaltered from Hay et al. (2011). *m* and *h* remain unitless gating variables in both equations. The *shift* parameters have units of mV.

Values of the maximum conductances associated with each of these currents in the Hay model and in our L5 Human model are given in Table 1.

**Table 1.** Parameters for the L5 Human model, with maximum conductances and passive properties compared to the Hay model. Parameters are shown to two decimal places or significant figures, although we note that NEURON has no limits regarding significant figures and thus further precision is used in the actual model implementation.

| **Ionic Current** | **L5 Human Model**<br>**max conductance**<br>**(S/cm<sup>2</sup>)** | **Hay Model**<br>**max conductance**<br>**(S/cm<sup>2</sup>)** | **H-current**<br>**Parameter** | **L5 Human**<br>**Model Value** |
| --- | --- | --- | --- | --- |
| Na_Ta (soma) | 2.10 | 2.04 | a, ms | 23.45 |
| Na_Ta (apical) | 0.0010 | 0.0213 | b, ms | 0.22 |
| Nap_Et2 | 1.90e-06 | 0.00172 | c, ms | 1.31e-09 |
| K_Pst | 0.065 | 0.00223 | d, ms | 0.083 |
| SKv3_1 (soma) | 0.040 | 0.693 | f, ms | 1.50e-09 |
| SKv3_1 (apical) | 0.040 | 0.000261 | k | 8.05 |
| SK_E2 (soma) | 2.45e-09 | 0.0441 | vh, mV | -90.87 |
| SK_E2 (apical) | 2.45e-09 | 0.0012 | ehcn, mV | -49.85 |
| K_Tst | 2.00e-05 | 0.081 | **"Shift" Parameter** | **Value** |
| Ca_LVA (soma) | 0.0010 | 0.0034 | *shift<sub>Na_Ta</sub>*, mV | -5 |
| Ca_LVA (apical) | 0.0010 | 0.019 | *shift<sub>SKv3_1</sub>*, mV | -10 |
| Ca_HVA (soma) | 5.70e-09 | 9.92e-04 |  |  |
| Ca_HVA (apical) | 5.70e-09 | 0.00056 |  |  |
| Ih (soma, basilar) | 5.14e-05 | 2.00e-04 |  |  |
| Im | 2.00e-04 | 6.75e-05 |  |  |
| **Passive Property** | **L5 Human Model** | **Hay Model** |  |  |
| **Parameter** | **Value** | **Value** |  |  |
| Ra, ohm cm | 495.73 | 100 |  |  |
| e_pas, mV | -84.40 | -90 |  |  |
| cm (soma), uF/cm<sup>2</sup> | 1.00 | 1.00 |  |  |
| cm (apical), uF/cm<sup>2</sup> | 1.60 | 2.00 |  |  |
| cm (basilar), uF/cm<sup>2</sup> | 1.60 | 2.00 |  |  |
| cm (axonal), uF/cm<sup>2</sup> | 1.60 | 1.00 |  |  |
| g_pas (soma), nS/cm<sup>2</sup> | 1.75e-05 | 3.38e-05 |  |  |
| g_pas (apical), nS/cm<sup>2</sup> | 1.75e-05 | 5.89e-05 |  |  |
| g_pas (basilar), nS/cm<sup>2</sup> | 1.75e-05 | 4.67e-05 |  |  |
| g_pas (axonal), nS/cm<sup>2</sup> | 1.75e-05 | 3.25e-05 |  |  |

#### Ion channel distributions

The locations of each of the 10 ion channel types in our human L5 pyramidal cell model are summarized in Table 2, and utilize a classification of each compartment in the neuron model as part of the soma, axon, or apical or basilar dendrites. With three exceptions, the ion channels were distributed as in the model of Hay et al. (2011).

**Table 2.**
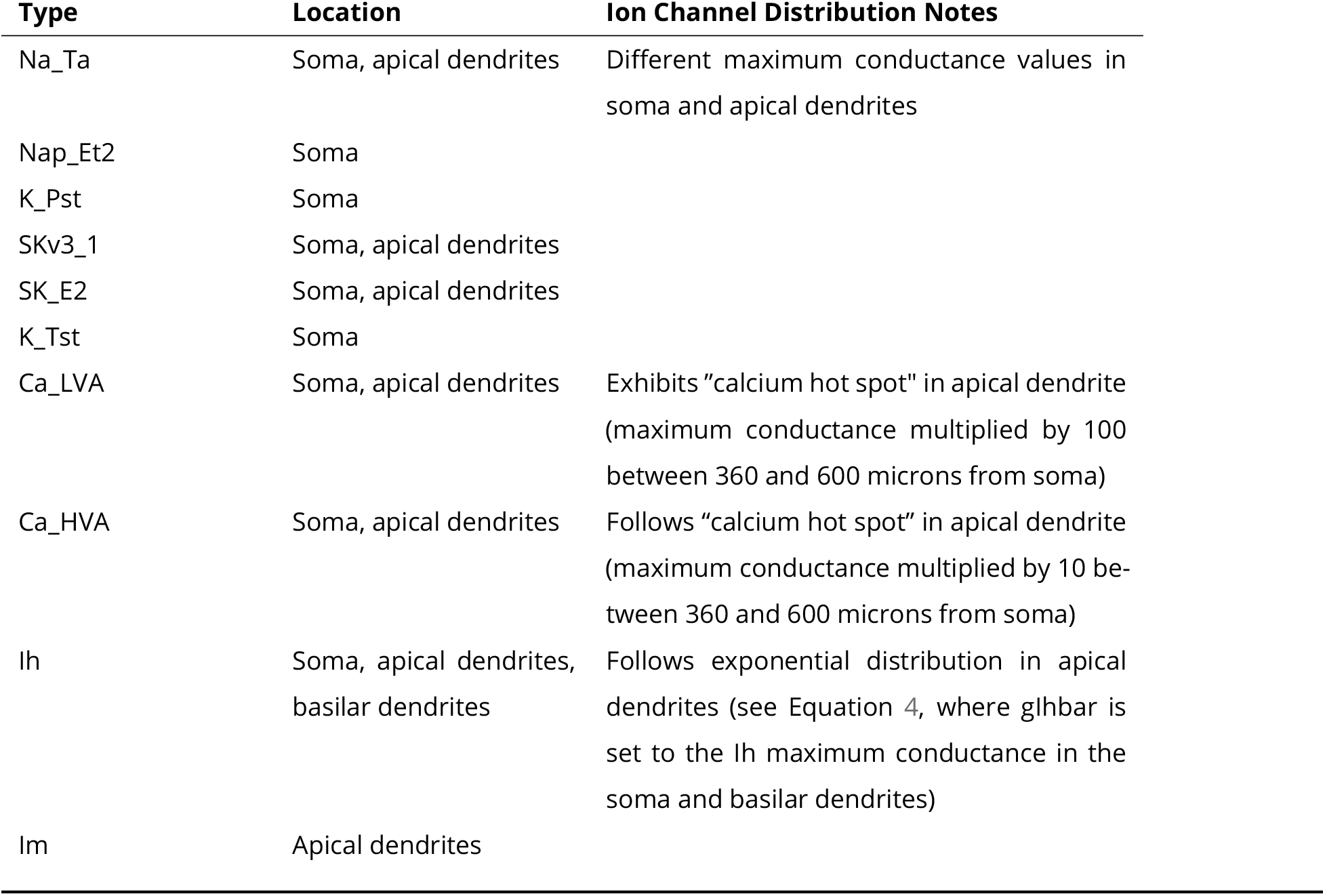
Summary of the distribution of ion channels in the differently classified compartments in the human L5 cortical pyramidal cell model.

| Type | Location | Ion Channel Distribution Notes |
| --- | --- | --- |
| Na_Ta | Soma, apical dendrites | Different maximum conductance values in soma and apical dendrites |
| Nap_Et2 | Soma |  |
| K_Pst | Soma |  |
| SKv3_1 | Soma, apical dendrites |  |
| SK_E2 | Soma, apical dendrites |  |
| K_Tst | Soma |  |
| Ca_LVA | Soma, apical dendrites | Exhibits "calcium hot spot" in apical dendrite (maximum conductance multiplied by 100 between 360 and 600 microns from soma) |
| Ca_HVA | Soma, apical dendrites | Follows "calcium hot spot" in apical dendrite (maximum conductance multiplied by 10 between 360 and 600 microns from soma) |
| Ih | Soma, apical dendrites, basilar dendrites | Follows exponential distribution in apical dendrites (see Equation 4, where $g_{lhbar}$ is set to the Ih maximum conductance in the soma and basilar dendrites) |
| Im | Apical dendrites |  |

The first and second exceptions are the calcium channels (Ca_HVA and Ca_LVA currents). A feature of the Hay et al. (2011) model that required adjustment was the “calcium hot spot”. As described by Hay et al. (2011) and Larkum and Zhu (2002), experimental evidence suggests a region of increased calcium channel conductance near the “main bifurcation” in the apical dendrites in rodent L5 pyramidal cells. The location of this bifurcation is closer to the soma in the morphology of the human L5 pyramidal cell than that used in Hay et al. (2011) considering the difference between human and rodent cell morphology, even in similar brain regions (Beaulieu-Laroche et al., 2018). As such the region of this increased calcium activity, where the Ca_LVA maximum conductance is multiplied by 100 and the Ca_HVA maximum conductance is multiplied by 10, is chosen to be on the apical dendrite 360 to 600 microns from the soma.

The third exception are the h-channels. The function used to model the “exponential distribution” of h-channels along the dendrites (Kole et al., 2006; Ramaswamy and Markram, 2015; Beaulieu-Laroche et al., 2018) was also slightly adjusted from that presented in Hay et al. (2011) given the distinct neuron morphology of the primary cell used here. For a given apical dendritic compartment, the maximum conductance of the h-current, gIhbar*, is given by the following equation:

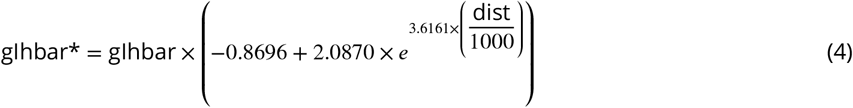

where “dist” is the distance from the soma to the midway point of the given compartment, the denominator of “1000” is chosen since this is the approximate distance from the soma to the most distal dendrite, and “gIhbar” is the h-current maximum conductance value that is optimized. “gIhbar” is also the value of the maximum conductance in the soma and basilar dendrites (i.e. the Ih maximum conductance is constant across all compartments in these regions).

### Details of the cycling fitting strategy

#### Parameter optimization using NEURON’s Multiple Run Fitter algorithm

The first step in the “cycling” model development strategy (schematized in Figure 2**A**) utilized NEURON’s built in Multiple Run Fitter (MRF) algorithm for optimization (Hines and Carnevale, 2001; Carnevale and Hines, 2006). This algorithm utilizes the PRAXIS method to minimize the mean squared error (MSE) between the output (in this case, a voltage trace) of the model neuron in comparison to experimental data obtained from an analogous protocol (Brent, 1976). Here, we fit the model to five different current clamp protocols experimentally obtained from the primary neuron from which we obtained our human L5 cell morphology (see Figure 1). As the experimental current clamp data was obtained in the presence of TTX, all sodium conductances were set to zero and not altered in this step. Additionally, the potassium channel currents primarily involved in action potential generation, K_Pst and SKv3_1, were omitted from the optimization and remained “set” during this step, as were the “shifts” associated with these sodium and potassium channels. Thus, the conductances that were determined in this first step were the SK_E2, K_Tst, Ca_LVA, Ca_HVA, Ih, and Im maximum conductances, along with the passive properties (in NEURON parlance Ra, e_pas, cm, and g_pas) and the kinetics of the h-current (in NEURON parlance a, b, c, d, f, k, vh, and ehcn).

**Figure 2.**
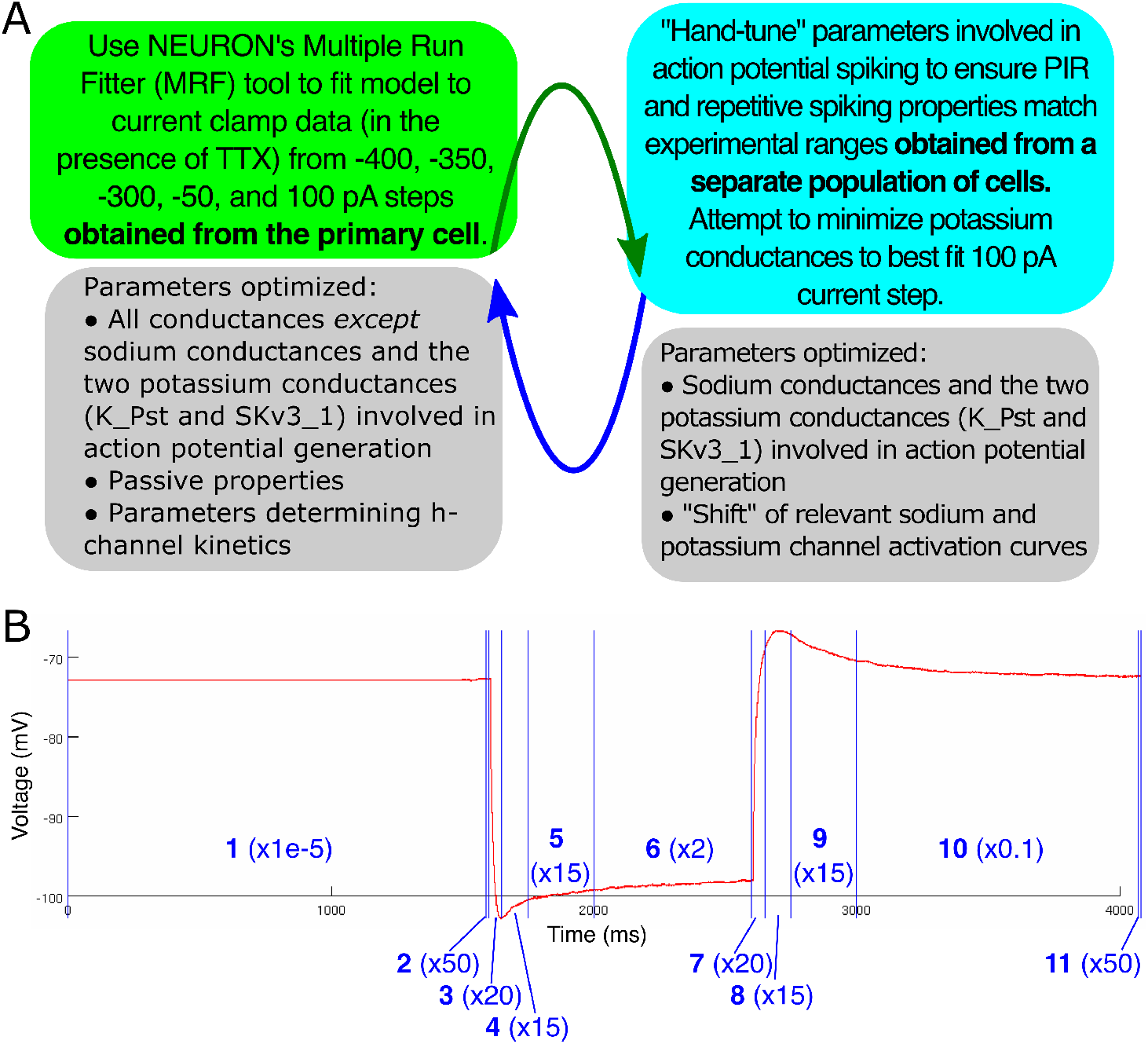
Details of model development strategy. **(A)** Diagram of the “cycling” modeling strategy. Hyperpolarizing current steps taken from the primary human L5 pyramidal cell were the primary constraint in determining model parameters. To ensure that the model exhibited repetitive and post-inhibitory rebound firing dynamics characteristic of human L5 pyramidal cells, data from a separate population of L5 cells was used to dictate a “reasonable range” of activity amongst human L5 pyramidal cells, and a “cycling” technique was developed in which conductances primarily active during spiking dynamics were adjusted by hand to ensure reasonable firing properties. The adjustments to the potassium conductances affect the current clamp fits, so these were re-run with the new values, hence the “cycle”. We note that, despite the use of data from the separate neuron population to inform the implementation of spiking characteristics in our model, only a single neuron model was generated, that using the “primary” neuron’s morphology. **(B)** Illustration of the differential “weighting” of the portions of the current clamp traces during optimization, used in the first step of the “cycling” strategy. The example experimental voltage trace shown here (a −400 pA step) *is* corrected for the liquid junction potential, as the model is directly fit to it. Each portion of the trace is numbered in blue, with the weight given to the error in this region included in parentheses. The highest weights are given to brief periods to match the experimentally observed resting membrane potential (2 and 11). The next highest weights are for the “charging” and “discharging” portion of the curve to emphasize the fitting of passive properties (3 and 7). The next highest weights are for the “sag” and “hump” most often associated with dynamics of the h-current (4, 5, 8 and 9). Portions of the curve associated with more “steady-state” behaviors are weighed significantly less given the aims of this modelling study (1, 6 and 10).

We chose to use three hyperpolarizing current clamp traces, with −400 pA, −350 pA, and −300 pA current amplitudes, because at these hyperpolarized voltages it was reasonable to assume that the h-current was primarily responsible for the voltage changes (Toledo-Rodriguez et al., 2004). This allowed us to accurately fit not only the h-current maximum conductance, but also its kinetics (see Equation 1 above).

A hyperpolarizing current step with a small (−50 pA) magnitude was chosen to constrain the passive properties, as near the resting membrane potential it is primarily these properties that dictate the voltage responses (“charging” and “discharging”) to a current clamp protocol. We note that this trace does not represent a perfectly “passive” neuron, as some conductances (such as those due to the h-current) are active, albeit minimally, at mildly hyperpolarized voltages (only the sodium channels were directly blocked in this protocol, via the application of TTX). Nonetheless, given that our model fit this current clamp data well, and also mimicked the “charging” and “discharging” portions of all the current clamp protocols included in the optimization, we are confident that we accurately approximated the passive properties of our particular human L5 pyramidal neuron. The final passive properties are shown in Table 1 along with those of a rodent L5 cortical pyramidal cell model of Hay et al. (2011). The passive properties include Ra (the axial resistivity in ohm cm), e_pas (the passive reversal potential in mV), cm (the specific capacitance in uF/cm^2^), and g_pas (the passive conductance in S/cm^2^).

Finally, a depolarizing current step (100 pA) was chosen to ensure the model was not “overfit” to the hyperpolarized data. Early in the modeling process, we recognized that a “best fit” of the depolarizing current clamp data would involve minimizing the values of the K_Pst and SKv3_1 maximum conductances to the point that action potential generation would not be viable. This was a critical motivation for the development of this “cycling technique” to ensure that reasonable spiking characteristics were achieved by the model while also minimizing these conductances as much as possible to best fit the depolarizing current clamp trace, and is discussed further below.

We note that, in the process of designing this modeling technique, we chose not to use every current clamp recording available to us, but instead chose a moderate number of current clamp recordings for use in the optimization. This allowed us to use the additional current clamp steps as “tests” not used in the fitting.

A useful tool provided by NEURON’s MRF is the ability to differentially “weigh” portions of the traces in the computation of the mean squared error value we sought to minimize. Given the focus of this study was on uncovering dynamics of the h-current, we more heavily weighed the portions of the voltage trace in which this channel most affected the voltage, namely the initial “sag” following a hyperpolarizing current steps and the “rebound” in voltage when this inhibition is released. We also chose portions of the voltage trace to emphasize in the mean squared error calculation in order to ensure the model cell closely approximated the resting membrane potential observed experimentally, as well as matched the “charging” and “discharging” features heavily influenced by passive properties. Where we report values of the MSE (particularly the captions of Figures 3 and 5) they reflect these weighting choices. We note that these differential “weights” were chosen only after a rigorous exploration of how these choices affected the overall model fit; indeed, this choice yielded a model that both qualitatively and quantitatively best fit the experimentally-observed behavior of our human L5 cortical pyramidal cell. These weighting choices are visualized in Figure 2**B**.

**Figure 3.**
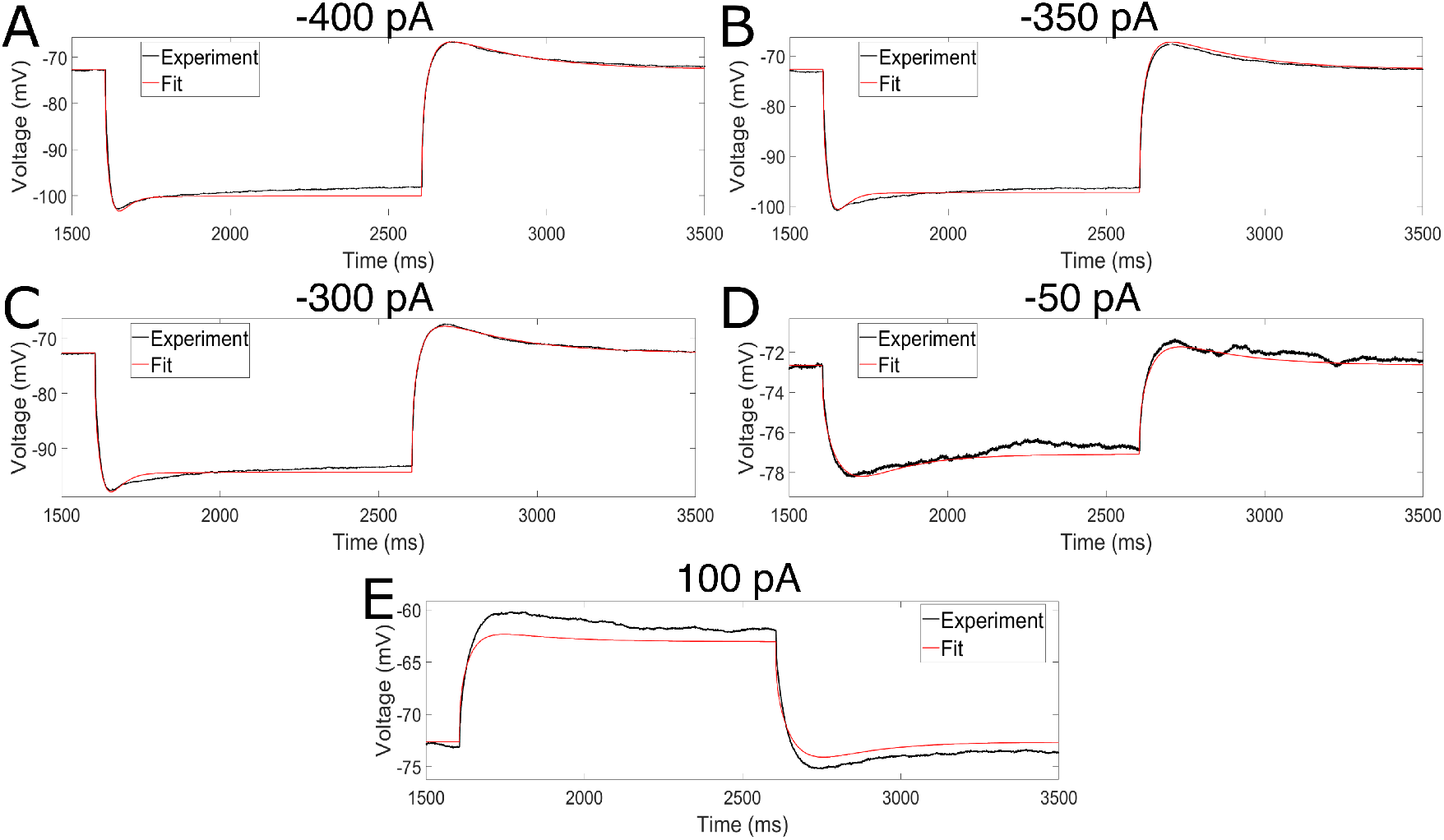
Model well fits data from hyperpolarizing current steps, in which the h-current is the primary active channel, while minimizing the error seen in a depolarizing current step. **(A-D)** Fits of hyperpolarizing current steps with −400 pA (MRF ouputted MSE of 0.194) **(A)**, −350 pA (MRF ouputted MSE of 0.309) **(B)**, −300 pA (MRF ouputted MSE of 0.130) **(C)** and −50 pA (MRF ouputted MSE of 0.027) **(D)** with TTX. **(E)** Fit of a depolarizing current step of 100 pA with TTX (MRF ouputted MSE of 1.029). All four hyperpolarizing current steps are fit with great accuracy, with a focus on the initial “sag” and post-inhibitory “rebound” that are driven by the activity of the h-current. While the charging and discharging portion of the depolarizing current trace is well fit, the amplitude of the response is less accurate; however, this error was deemed reasonable given the emphasis in model development on capturing h-current dynamics, including PIR spiking, as discussed in detail in the text.

**Figure 4.**
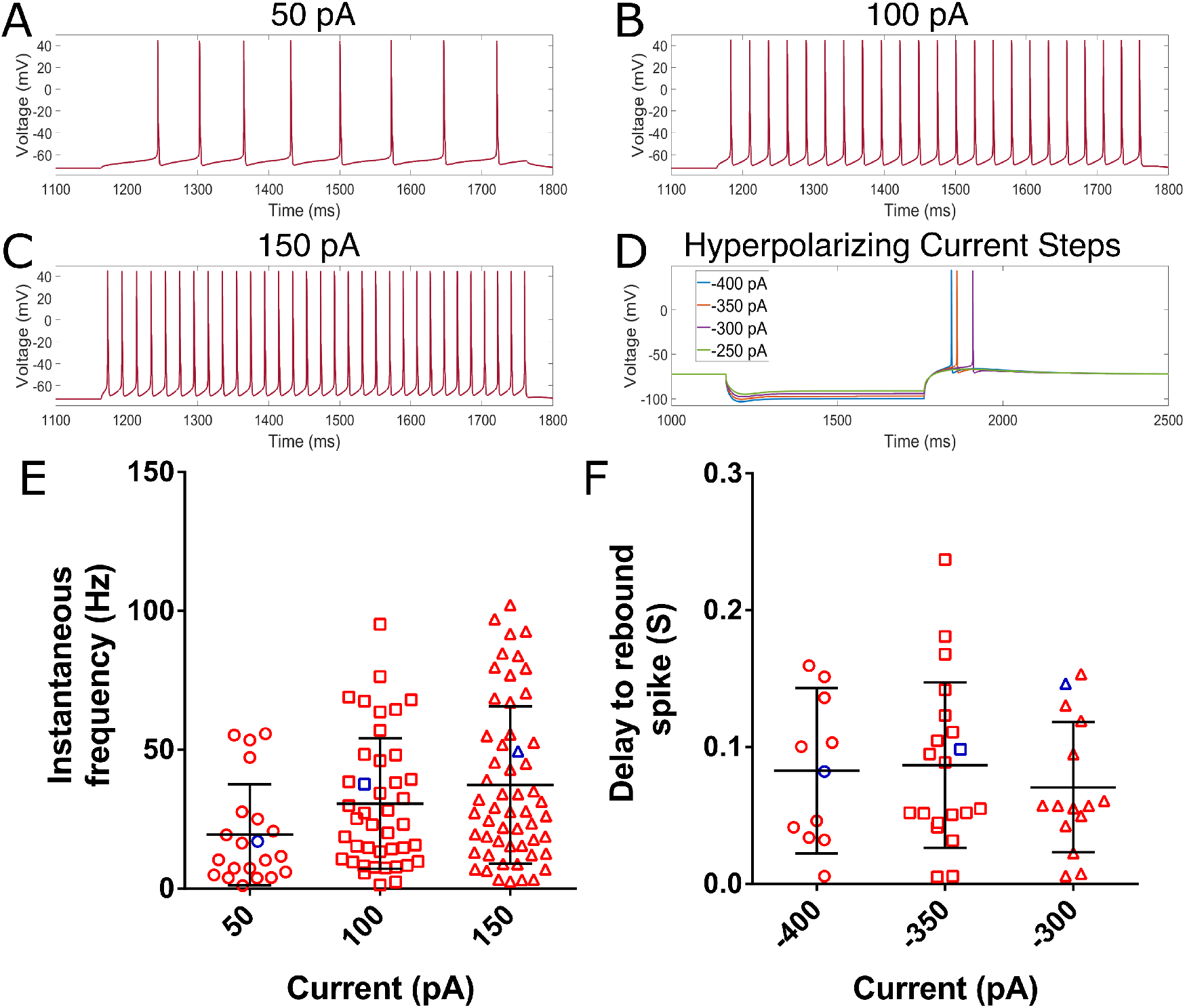
Model neuron exhibits reasonable repetitive and PIR spiking behavior. **(A-C)** Repetitive spiking behavior of the model neuron in response to 50 pA **(A)**, 100 pA **(B)**, and 150 pA **(C)** current steps. **(D)** PIR behavior in response to four hyperpolarizing current steps. **(E)** Butterfly plots of the instantaneous frequency (i.e. the frequency derived from the cell’s first two spikes) of a population of human L5 pyramidal cells characterized by Chameh et al. (2019), with black bars representing the mean ± standard deviation. The blue marker represents the value for our model neuron. **(F)** Analogous plots to panel **E** from the same cells, but for the delay to the PIR spike following the release of the hyperpolarizing current step. Only cells that exhibit a PIR spike are shown here (as discussed in the Methods). In both panels **E** and **F** we see that the value derived from our model neuron (again, in blue) falls within the range exhibited by these neurons experimentally, often quite near the average value.

**Figure 5.**
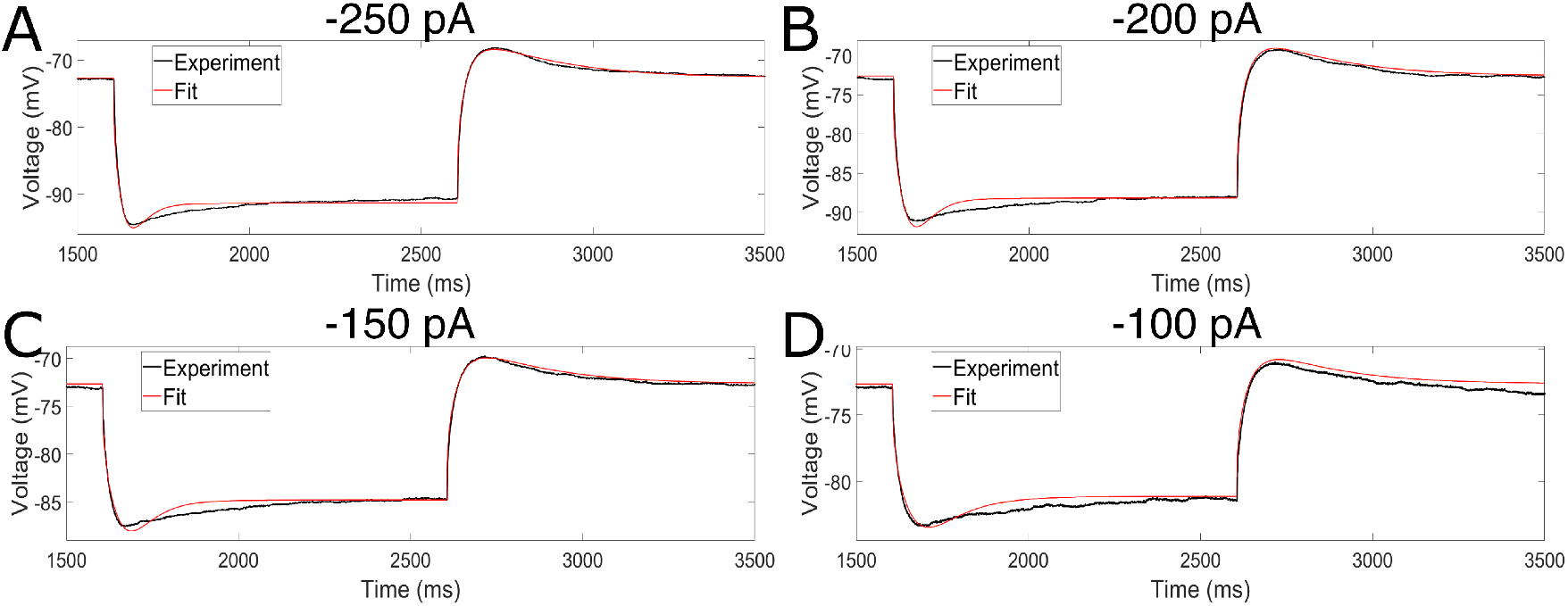
Model output well-matches experimental data not used as constraints in model generation. **(A-D)** Voltage traces from current clamp experiments with TTX for steps of −250 pA (MSE with MRF weights of 0.145) **(A)**, −200 pA (MSE with MRF weights of 0.275) **(B)**, −150 pA (MSE with MRF weights of 0.197) **(C)** and −100 pA (MSE with MRF weights of 0.329) **(D)**. Model output (red curve) well matches the experimental observations (black curve), with MSE errors of similar magnitude to the traces directly fit to (see caption of Figure 3) despite these particular traces not being used in model generation.

**Figure 6.**
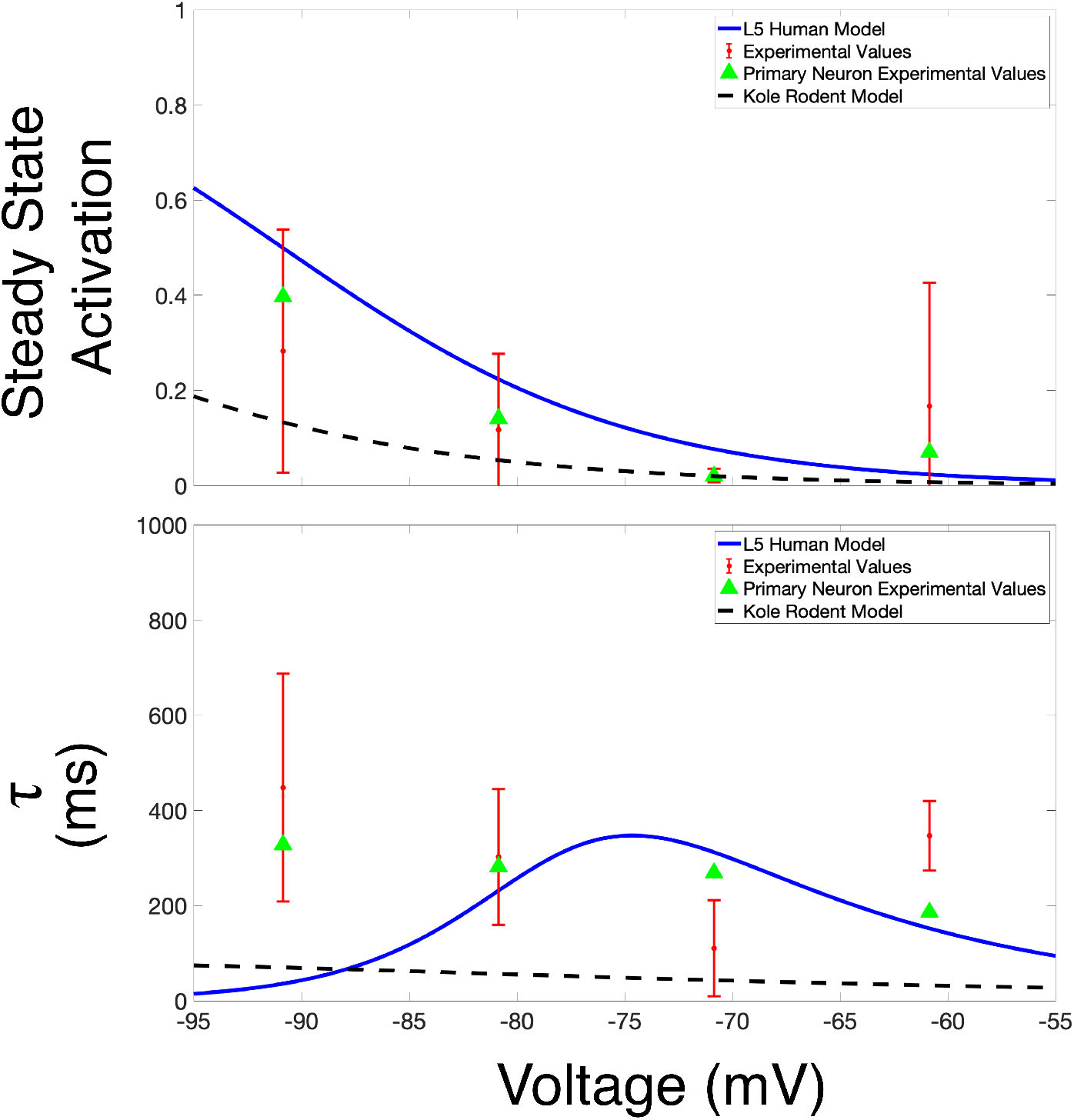
The human h-current model presented in this work is validated by comparison to experimental voltage clamp data. Plots of steady-state activation (top row) and *r* (bottom rows) curves over an illustrative voltage range (discussed in more detail in the text). The values for our L5 human h-current model are shown in blue, with these values juxtaposed with those extracted from voltage clamp experiments: data from the primary cell are shown via green triangles, and data averaged over a secondary population L5 cortical pyramidal cells (with the standard deviation shown via error bars) are shown in red. For comparison, analogous curves from the Kole et al. (2006) rodent h-current model are shown via a dotted black line.

**Figure 7.**
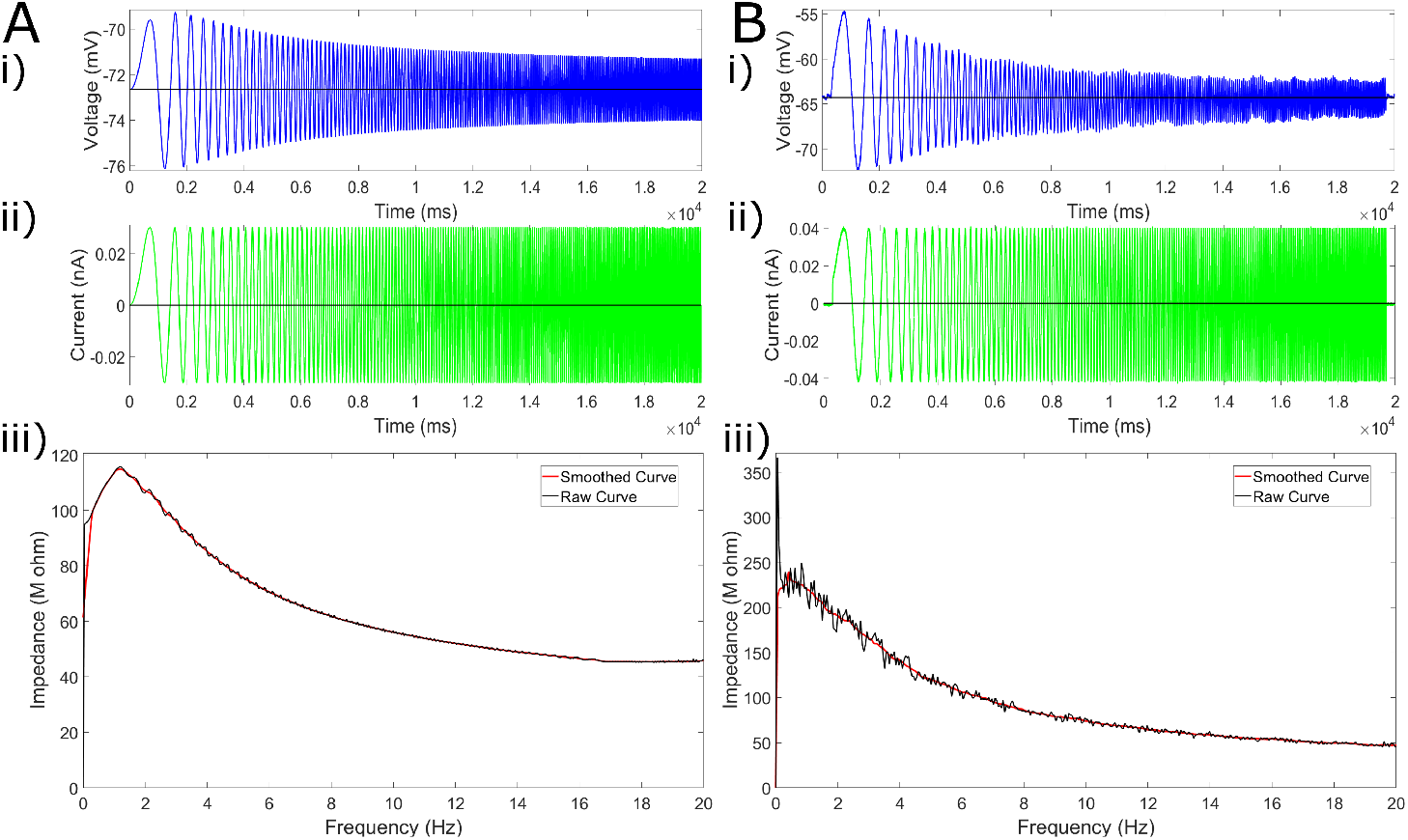
Model matches experimental data from human L5 pyramidal neurons lacking subthreshold theta resonance in response to ZAP function input. **(A)** *In silico* results from the model neuron to subthreshold current input from a ZAP function. The voltage response is shown in **i)**, the input current in **ii)**, and the calculated impedance in **iii)**, illustrating the lack of a peak at theta frequency. **(B)** Example *in vitro* results of an analogous ZAP protocol (plots correspond with those in panel **A**) show the lack of subthreshold resonance experimentally (see Chameh et al. (2019)).

We further note that the possibility that our parameters at the end of this step of model generation represented a “local” rather than “global” minimum in the optimization was considered and thoroughly tested to confirm the robustness of this process. This included running the optimization with a variety of initial conditions, as well as “perturbing” individual parameters to see whether the optimization would converge back; this was performed not only with the final model, but at intermediate stages in its development. We thus are quite confident that the model presented in this work represents a “global minimum” of the MRF algorithm as implemented here.

#### Matching of spiking features

In the second step of our model generation process, we tuned the sodium and potassium conductances involved in action potential generation (Na_Ta and Nap_Et2 sodium maximum conductances and the K_Pst and SKv3_1 potassium maximum conductances) by hand. The two “shift” parameters (*shift*_*Na*_*Ta*_ and *shift*_*SKv*3_1_) were also hand-tuned in this step. The primary goal of this step was to ensure the model exhibited PIR spiking, considering this dynamic was shown by Chameh et al. (2019) to be dependent upon the activity of the h-current in human L5 pyramidal neurons. In parallel, we sought to have our model neuron fall within a reasonable “range” of spiking properties observed in the separate population of human L5 pyramidal cells from Chameh et al. (2019), illustrated in Figure 4**E** and **F**.

We aimed to obtain these firing characteristics with minimal potassium conductances, in order to minimize the error seen in Figure 3**E**: an extensive exploration of the parameter space revealed that a “best fit” of this trace would enforce values of the potassium conductances that would not permit action potential firing, motivating the hand tuning of these values in search of a set of sodium and potassium conductance values that would permit spiking while also minimizing this error. Imparting reasonable spiking characteristics onto the neuron allows this model to be used as a “backbone” for future computational studies into suprathreshold behaviors such as frequency-dependent gain (a measure described by Higgs and Spain (2009) and used by Chameh et al. (2019) to characterize L5 pyramidal neurons *in vitro*).

In this step, we also found that a “shift” in the activation curve for Na_Ta (see Equation 2 above) was necessary to achieve PIR spiking as commonly seen experimentally. We sought to minimize this shift because a side effect of this leftward shift was an increase in repetitive firing frequency that approached the upper limit of what was biologically reasonable. We note that the final shift of −5 mV kept the dynamics of our sodium channel well within a reasonable range (for example, the sodium channel used in the model presented by Ascoli et al. (2010) has a significantly more leftward shifted sodium activation curve than our model).

Finally, in order to prevent biologically unrealistic depolarization blocks from occurring in our model (since these are not seen experimentally), we shifted the activation curve for SKv3_1 more leftward (−10 mV) than the sodium channel (see Equation 3 above). This technique for preventing depolarization block in computational models (i.e. shifting potassium current more leftward than the sodium current) has been previously suggested by Bianchi et al. (2012).

We note that the features altered during this step were not altered in the preceding optimization step, both due to the presence of TTX in the current clamp recordings and in order to ensure the ability of our neuron to spike was not compromised by “overfitting” non-spiking data.

#### Final model parameters

As the properties of the potassium channels altered in the pursuit of matching spiking features affected the current clamp fits, it was then necessary to run the optimization algorithm of the first step again with these new values; hence the “cycling” fitting strategy. The “cycling” mechanism was run until there was no significant improvement in the quantitative (i.e. the mean squared error outputted by the MRF algorithm in the optimization step) or qualitative (i.e. the spiking characteristics) measurement of model accuracy in either step of the cycle. The resulting parameter choices are summarized in Table 1, shown together with those of a rodent L5 pyramidal cell as developed by Hay et al. (2011).

The input resistance of the final model was 82.16 Mohm which compares favourably with the experimental data from the primary cell which yields an input resistance of 82.08 Mohm. This correspondence is as expected given the accurate fits that drove the modeling process. These values were determined by performing a linear fit (with a fixed y-intercept of 0) between an input current (“x value”) and the resulting steady-state change in voltage (“y value”) for input currents of −200, −150, −100, −50, 0, 50, and 100 pA (with sodium channels blocked, as in the experimental setting).

We obtain an approximation of the membrane time constant of both our model and the experimental neuron by fitting a double-exponential equation (*a* * *e*^*b*\**x*^ + *c* * *e*^*d*\**x*^) to the discharging portion of the voltage trace in response to the −50 pA current clamp, with the membrane time constant being the inverse of the constant corresponding with the “slow” exponent (i.e. the value of *b* or *d* that was smaller in magnitude). The membrane time constant of our final model was 27.53 ms, which compares favourably with the experimental data from the primary cell which yields a membrane time constant of 30.60 ms. Again, this correspondence is as expected given the accurate fits that drove the modeling process. We note that the time constant as calculated here is not the exact “RC” time constant, but rather an approximation of the neuronal membrane’s activity as commonly captured experimentally (see, for example, the calculation of the time constant presented by Chameh et al. (2019)).

Finally, we note that an approximation of the reversal potential of the h-current can be derived from voltage clamp data. While we were unable to obtain the full suite of voltage clamp data necessary for this calculation in our primary cell, we were able to do so in some of our secondary population of L5 cortical pyramidal neurons. This experimentally derived reversal potential should be considered alongside the caveats that must be considered with the voltage clamp data (discussed at length above), but is still likely a close approximation of the actual value of this parameter. Using three cells, each cell was voltage clamped at −40 mV (values not corrected for the liquid junction potential), and stepped down in 10 mV increments. The amplitude of the h-current was measured at each voltage, and a fit line was extrapolated to find the voltage at which the amplitude of the h-current would be 0. This was found to be −39.46 mV, which when corrected for the liquid junction potential is −50.26 mV. This value is a near perfect match to the value obtained via our modeling process of −49.85 mV. This can be viewed as further validation that our modeling process reasonably approximated the biological reality of the h-current in human L5 pyramidal neurons.

#### Parameter constraints

Moderate constraints were placed on the range of certain parameters in order to ensure that, in finding the best “fit” to the data, these values did not enter a regime known to be biologically unlikely or that would lead to unreasonable spiking characteristics. In order to preserve reasonable spiking behavior, the maximum value for the Ca_LVA maximum conductance was set to 0.001 S/cm^2^, the maximum value for the Ca_HVA maximum conductance was set to 1e-05 S/cm^2^, and the minimum value of the Im maximum conductance was set to 0.0002 S/cm^2^. These values were determined after rigorous investigation of the effects of these maximum conductances on the spiking properties.

Further constraints were placed on the passive properties of the neuron to make sure the neuron not only matched “charging” and “discharging” properties in the current clamp data, but also reasonably approximated the resistance and membrane time constant values generally seen in the population of L5 neurons studied by Chameh et al. (2019). We note that the variability in these values are higher than typically seen in the rodent setting, although this increased variability in such properties in human neurons is seen in a variety of recent studies (Chameh et al., 2019; Kalmbach et al., 2018; Beaulieu-Laroche et al., 2018; Chartrand et al., 2019; Gidon et al., 2020), which justified our decision to not overly constrain these parameters. These limits were as follows: the axial resistance (Ra) was constrained between 0 and 1000 ohm cm; the membrane capacitance (cm) outside the soma was constrained between 1 and 1.8 uF/cm^2^; the passive reversal potential (e_pas) was constrained between −90 and −80 mV; and the passive conductance (g_pas) was constrained between 1.75e-05 and 2.5e-05 S/cm^2^.

### *In silico* experiments

The usefulness of the model presented here lies not only in its ability to well “fit” the constraining data, but the insights it provides when subjected to *in silico* versions of experiments. A common protocol used to assess sub-threshold neural activity was performed *in silico* on our model neuron to evaluate the ability of our neuron model to capture an “essence” of the functional capacity of the neuron, and this data was compared to available results from analogous *in vitro* experiments.

A “ZAP function”, a sinusoidal function whose frequency changes linearly over a given range, has been used to assess the impedance amplitude profile in a variety of engineering settings for over 30 years (Puil et al., 1986), including in the assessment of subthreshold resonance properties in neurons (Leung and Yu, 1998). In this study, the ZAP function protocol was motivated by that used in the corresponding experimental data (Chameh et al., 2019): the current injection lasted for 20 seconds with its frequency ranging from 0 to 20 Hz. The current was injected into the soma of the model, just as the experimental protocol was somatic. The amplitude of this input was 0.03 pA in all *in silico* protocols. The ZAP current was delivered after a delay in order to allow the neuron to equilibrate at its resting membrane potential (or the steady-state potential given a particular DC current) before ZAP application.

We note that, in Figure 7**A**, only a single experiment is shown. As the ZAP current is set and the model neuron is deterministic (i.e. will exhibit the same response to the same input in every case), no averaging or statistical measures were necessary for this protocol. We also note that, in determining the “resonant frequency” highlighted in Tables 4, 5, and 6, we only consider frequencies greater than 1 Hz. The peak values displayed in these tables were found simply by determining the frequency corresponding to the maximum impedance value (in the raw, rather than “smoothed”, data).

The code generating this current was obtained from the NEURON (Carnevale and Hines, 2006) website via the following link: http://www.neuron.yale.edu/ftp/ted/neuron/izap.zip.

### Implementation of other models

Models from two other works, that of Hay et al. (2011) and Kalmbach et al. (2018), were implemented and used for comparison purposes.

The Hay et al. (2011) model is accessible via ModelDB at senselab.med.yale.edu/ModelDB (Accession:139653). We implemented this model directly using the code available via this source. In this work we utilized the model that is “constrained both for BAC firing and Current Step Firing”, which is dictated by specifically utilizing the “L5PCbiophys3.hoc” file.

The Kalmbach et al. (2018) model is available via GitHub at https://github.com/AllenInstitute/human_neuron_Ih. The morphology of the model neuron and the “shifted” version of the Kole et al. (2006) h-current model that are used were directly downloaded from this repository, and the passive properties and h-current maximum conductance values as defined in the code repository were instantiated via basic NEURON code. This “shifted” version of the Kole et al. (2006) model is included below:

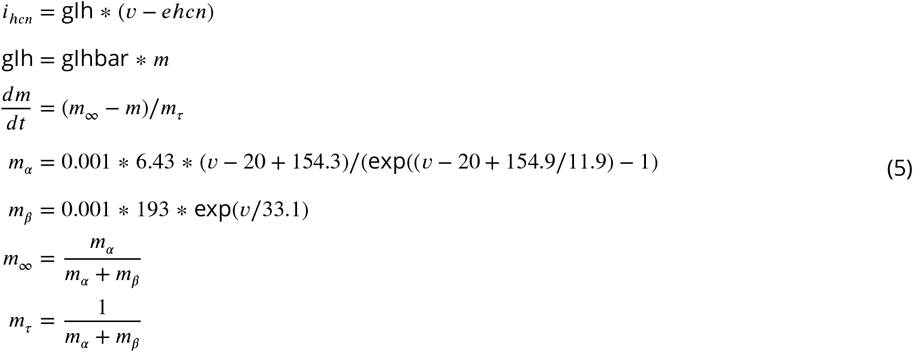

The “-20” term in the *m*_*α*_ equation is the “shift” from Kole et al. (2006). The parameters dictating the model which has non-uniform passive properties and uniformly distributed h-channels (amongst the soma, apical, and basilar dendrites) are given in Table 3. We ensured our implementation of this model was appropriate by directly replicating Figure 7B of Kalmbach et al. (2018) with this implementation.

**Table 3.**
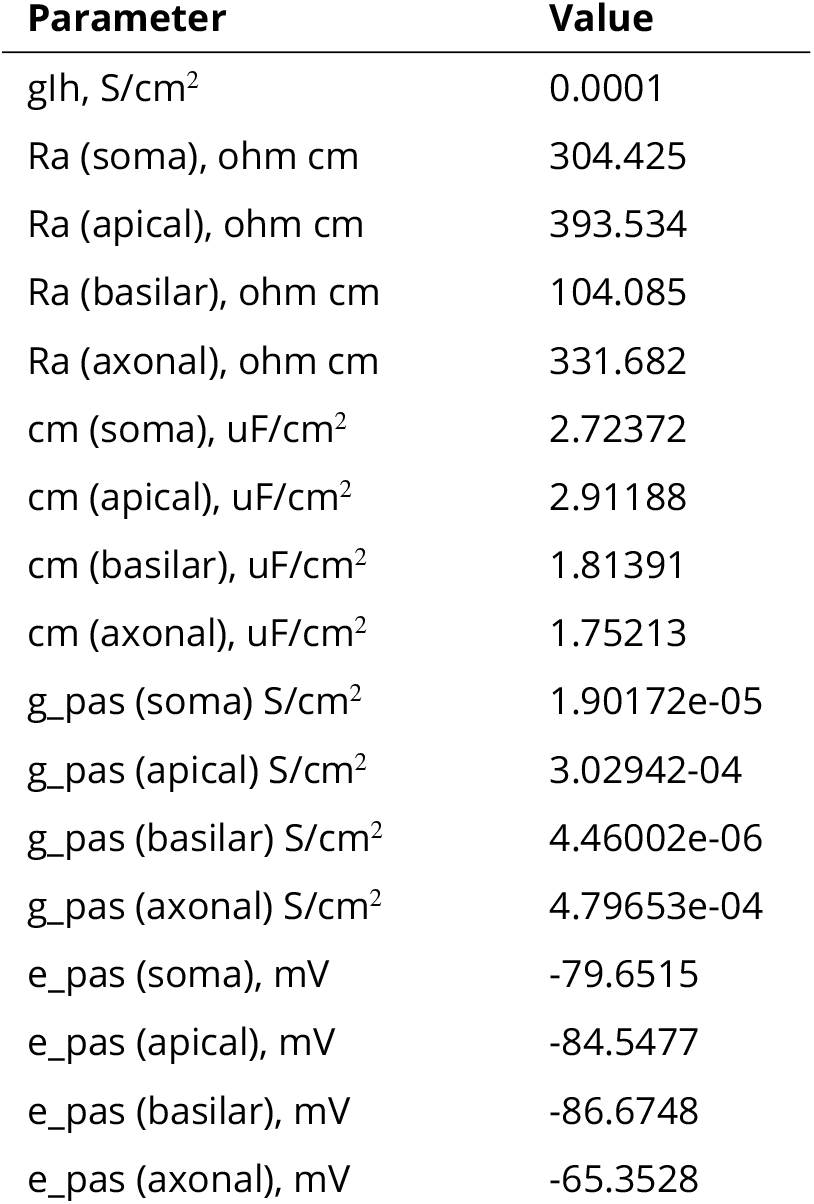
Exact parameters used in implementation of the human L3 cortical pyramidal cell model of Kalmbach et al. (2018).

**Table 4.** Quantified results of the ZAP protocol applied to the three pyramidal cell models of interest highlight different propensities for subthreshold resonance.

| Model | Conditions | RMP | Frequency of Peak Impedance (>1 Hz) |
| --- | --- | --- | --- |
| L5 Human | Default | -72.43 mV | 1.19 Hz |
| L5 Human | Block all channels besides h-channel | -72.01 mV | 1.22 Hz |
| Hay | Default | -77.25 mV | 2.80 Hz |
| Hay | Block all channels besides h-channel | -76.87 mV | 2.79 Hz |
| Kalmbach | Default (model's only active ion channel is the h-channel) | -78.41 mV | 4.00 Hz |

**Table 5.** Quantified results of the ZAP protocol applied with DC shifts to the three pyramidal cell models of interest (with all included ionic currents active)

| Model | DC shift | RMP | Frequency of Peak Impedance (>1 Hz) |
| --- | --- | --- | --- |
| L5 Human | -100 pA | -81.11 mV | 2.08 Hz |
| L5 Human | -200 pA | -88.16 mV | 2.66 Hz |
| L5 Human | -300 pA | -94.31 mV | 3.03 Hz |
| L5 Human | -400 pA | -100.05 mV | 3.71 Hz |
| Hay | -100 pA | -81.04 mV | 3.07 Hz |
| Hay | -200 pA | -84.47 mV | 3.13 Hz |
| Hay | -300 pA | -87.55 mV | 2.99 Hz |
| Hay | -400 pA | -90.28 mV | 3.91 Hz |
| Kalmbach | -100pA | -80.28 mV | 4.42 Hz |
| Kalmbach | -200pA | -82.05 mV | 4.43 Hz |
| Kalmbach | -300pA | -83.73 mV | 5.06 Hz |
| Kalmbach | -400pA | -85.34 mV | 4.83 Hz |

**Table 6.** Quantified results of the ZAP protocol applied to “hybrid” models with and without DC shifts

| Model “back-bone”: | H-current model: | Name of “hybrid” model | DC shift | RMP | Frequency of Peak Impedance (>1 Hz) |
| --- | --- | --- | --- | --- | --- |
| Hay | L5 Human | Hay-L5 Human hybrid | 0 pA | -72.43 mV | 1.16 Hz |
| Hay | L5 Human | Hay-L5 Human hybrid | -200 pA | -77.95 mV | 2.20 Hz |
| Hay | L5 Human | Hay-L5 Human hybrid | -400 pA | -82.41 mV | 2.20 Hz |
| Kalmbach | L5 Human | Kalmbach-L5 Human hybrid | 0 pA | -78.37 mV | 2.28 Hz |
| Kalmbach | L5 Human | Kalmbach-L5 Human hybrid | -200 pA | -81.72 mV | 3.03 Hz |
| Kalmbach | L5 Human | Kalmbach-L5 Human hybrid | -400 pA | -84.68 mV | 3.44 Hz |

In both cases, replacing the default rodent-motivated h-current model with the h-current model generated in this study was a straightforward matter of changing which channel was added into the NEURON model. Doing so ensured that the *only* change in these “hybrid” models was to the kinetics of the h-current (i.e the h-channel distribution and maximum conductance, as well as all other features, were the same as in the “model backbone”). All code involved in the implementations of these models is available at https://github.com/FKSkinnerLab/HumanL5NeuronModel.

## Results

### Development of a human L5 cortical pyramidal cell model using a cycling fitting strategy

In developing a detailed, biophysical model of a given cell type it is preferable to use data from the same cell, as averaging experimental data from multiple cells does not necessarily capture the particular cell’s characteristics and can be erroneous (Marder and Taylor, 2011). However, obtaining a full suite of data necessary to characterize the different ion channel types individually for a particular cell type is essentially impossible. This is additionally challenging when building human cellular models due to limited tissue access. Given these considerations, we developed a “cycling” fitting strategy to best utilize our unique human data set to build our model. Data from three sets of neurons were utilized: our primary neuron yielded a detailed morphology and electrophysiological recordings in the presence of TTX (the current clamp data from this neuron is shown in Figure 1); a suite of secondary neurons yielded voltage clamp data that characterized the h-current in the same fashion as done in the primary neuron; and a set of neurons first characterized in previous work (Chameh et al., 2019) yielded spiking characteristics considering they were not treated with TTX.

The “cycling” technique schematized in Figure 2**A** is described in detail in the Materials and Methods. Briefly recapped, we first use a built-in optimization technique in the NEURON (Carnevale and Hines, 2006; Hines and Carnevale, 2001) platform to best-fit current clamp data, and then “hand-tune” parameters primarily associated with action potential generation in order to ensure the neuron exhibited reasonable spiking characteristics. This multi-step process was necessary given the complex interactions between the various parameters influencing both the fit to current clamp data and spiking behavior. The resulting model not only facilitates the detailed investigation into the kinetics of the h-current and subthreshold resonance described below, but also allows for this model to serve as a “backbone” for future, more detailed investigations into spiking properties of distinctly human neurons. Some of these potential future applications will be elaborated on in the Discussion section.

The output of our final model in response to the various current steps with blocked sodium channels, compared to what was observed experimentally in the primary neuron, is shown in Figure 3**A-E**. The repetitive spiking behavior of the model in response to various driving currents is shown in Figure 4**A-C**, and the capacity for PIR spiking is shown in Figure 4**D**; both of these protocols are performed with active sodium channels. The instantaneous firing frequency (i.e. the frequency derived from the neuron’s first two spikes) or latency to the first PIR spike (depending upon whether the protocol is a depolarizing or hyperpolarizing current clamp, respectively), is compared to separate experimental data from Chameh et al. (2019) in Figure 4**E** and **F**.

Critically, the model closely matches this experimental data presented in Figure 3, indicating that the dynamics of the h-current within this voltage range were accurately encapsulated by our model. While the error in the depolarizing current step (Figure 3**E**) is more noticeable, this was minimized via the process described above, and was the best case while also ensuring reasonable repetitive spiking and PIR spiking behaviors (Figure 4**E** and **F**). Indeed, the spiking frequencies and latencies to the PIR spike highlighted in Figure 4**E** and **F** all fall within the range exhibited by the experimental data from our separate population of L5 pyramidal cells. Thus, this error in the depolarizing current step was deemed a necessary concession in order to preserve the accurate replication of h-current driven behaviors (current clamp fits to hyperpolarizing steps and the capacity for PIR) and to make a model that has the potential for more general application (given its reasonable spiking characteristics).

While the overall fits to the current clamp traces are convincing, there are discrepancies that are worth highlighting: specifically, while the initial “sag” voltage traditionally associated with h-current driven dynamics is well accounted for, this sag in our model tends to dissipate faster than in the experimental setting, and the model reaches a “steady-state” voltage much earlier than the biological cell. There are a variety of possible explanations for this discrepancy, all of which emphasize the fact that models are necessarily an abstraction of the biological entity and can not be expected to encapsulate every aspect of a neuron’s behavior (see the detailed analysis of this point in the Discussion); indeed, in the modeling process we made a conscious choice to create a model that captures the dynamics of the h-current as accurately as possible, sometimes at the expense of similar precision in other aspects of the model. This leads to one potential explanation for this discrepancy, which is that it is a result of the relative “weighting” of the errors in the MRF algorithm that emphasized fitting the initial sag (see Figure 2**B**). Another very likely explanation is that other ionic currents are at play in these dynamics that are not accounted for in the model; indeed, it is a near-certainty that the 10 ionic currents included in our modeling process do not fully encapsulate all of the currents in the biological neuron, especially considering these currents were chosen based on the rodent literature given the lack of similarly detailed human literature. The fact that our model better captures the speed at which the neuron returns to the resting membrane potential following the current clamp, when compared to how it captures the speed at which the neuron approaches a steady state during the current clamp, indicates that perhaps other ion channels not incorporated in this model contribute to the biological neuron’s behavior at hyperpolarized voltages. Focused experiments to understand what other currents might specifically contribute to the dynamics of human L5 pyramidal cells could thus lead to improvement of this model in the future.

We emphasize that we did not directly fit our model to the average values of these spiking properties, but instead aimed for these properties to fall in the experimentally observed “range”. This choice is motivated not only by the impact of cell-to-cell variability, but also the increased variability between similarly classified cells observed in humans in comparison to rodents (Chameh et al., 2019; Kalmbach et al., 2018; Beaulieu-Laroche et al., 2018; Chartrand et al., 2019; Gidon et al., 2020). Correspondingly, it would be inappropriate to directly fit or constrain the spiking properties of our model with these averages, or display such a comparison in Figure 4**A-D**. We also note that both the experimental and modeling data is intended to make no claims about rather PIR occurs in these cells under *in vivo* or more physiological settings; rather, the observed capacity for these neurons to exhibit this dynamic in *in vitro* settings is used as a tool for model development, especially considering the known complicity of the h-current in this behavior.

To justify our assertion that this model is appropriate for use in settings beyond those directly constraining model generation, we provide additional validating evidence in three capacities. This serves to rule out the possibilities that we either accidentally “overfit” our model to the chosen constraining data, or that this chosen data was somehow idiosyncratic and not indicative of the general properties and dynamics of the primary neuron and human L5 cortical pyramidal cells generally. First, we test the model against current clamp data obtained experimentally from the primary neuron but not used in model development (below); second, we compare the dynamics of the modeled human h-current to those observed experimentally both in the primary neuron and the secondary population (in the following section); and third, we compare the model’s capacity for subthreshold resonance with that observed experimentally in human L5 cortical pyramidal cells generally by Chameh et al. (2019) (in the following section).

Figure 5 illustrates the output of the model with four hyperpolarizing current steps, in comparison to the experimentally observed output from primary neuron, that were not directly “fit” in model generation. We again focus on hyperpolarizing current steps given the focus on the h-current, which is activated at hyperpolarized voltages, in this endeavor. The strong correspondence between the model and the experimental data illustrates that the modeling process described here does indeed capture the general behavior of the primary neuron with its morphological and passive property characterization in response to hyperpolarizing current steps of varying amplitudes. Perhaps most importantly, in all four cases the features of the trace most prominently influenced by the h-current, the initial “sag” following the onset of the hyperpolarizing current step and the “rebound” following its release, remain reasonably approximated by the model. We note that similar discrepancies between the model and experimental traces that were seen in Figure 3 persist in these examples, and further discrepancies in the magnitude of the “sag” can be seen; however, this is to be expected since these traces are “tests” of the model, rather than examples of model “fits”. Indeed, the fact that the error scores of these tests (included in the figure caption) are on the order of the errors seen in the fitting itself is convincing evidence that our model can capture the general features of the dynamics of a human L5 neuron beyond the specific setting in which it was constrained.

This result is a straightforward way of assessing our model’s validity via its ability to well match additional current clamp traces from the primary neuron. We note that this validity is driven not only by the focus on the h-current, but by fitting the other maximum conductances as well: as seen in Table 1, all the included maximum conductances differ from the values in the rodent L5 model of Hay et al. (2011), in some cases rather notably. Nonetheless, considering the h-current’s dominance over the neuron’s dynamics at hyperpolarized voltages, this result also provides more specific support for our assertion that our model captures the dynamics of the h-current.

### Model replicates experimental h-current kinetics and subthreshold resonance features

Allowing for distinct “human” kinetics of the h-current, relative to those observed in rodents (Kole et al., 2006), was paramount in facilitating the accurate fits of the *in silico* model (see Figure 3) to the *in vitro* experimental data presented in Figure 1. Such dynamics were constrained solely via the cycling optimization technique summarized above. With these fits in hand alongside the presence of additional experimental data, namely voltage clamp recordings from both the primary neuron and secondary human L5 cortical pyramidal cells (described in detail in the Materials and Methods), approximate experimental values of the voltage dependence of the time constant (denoted *τ*) and the steady-state activation values of the h-current are obtained and compared with the h-current model derived from our modeling process.

In Figure 6 we present experimental values of these quantities, alongside the human h-current model as well as the model of Kole et al. (2006) that was used by Hay et al. (2011). There are multiple reasons we would not expect our model to perfectly match this experimental data, including space-clamp issues associated with voltage clamp recordings, the fact that these recordings are somatic and h-channels are distributed throughout the dendrites in our model following rodent-like distributions, and the possibility that other ionic currents active at hyperpolarized voltages are not accounted for in our model and not blocked by our experimental protocol. Nonetheless, these experimental values can be considered as imperfect “approximations” of the biological kinetics of the h-current given our current understanding of the ionic channels active at hyperpolarized voltages, and are thus useful as a secondary validation that the kinetics derived in our modeling process are reasonable. This is especially important considering the notable difference between the kinetics of the h-current derived in this model and the rodent model of Kole et al. (2006); in particular, our model has values of *τ* nearly an order of magnitude larger, indicating the h-current acts in a significantly slower fashion. Additional justification that this is biologically reasonable is incredibly useful, despite the caveats that must be considered alongside the voltage clamp data.

Importantly, this experimental data *does* justify this human-rodent difference predicted by our modeling. Figure 6 shows that the human h-current model much more closely matches the *τ* values derived from the data from our primary neuron. While the superiority of either model is less conspicuous in the steady state activation values, the human model more closely matches the data from our primary cell at more hyperpolarized voltages at which the qualitative “shape” of this curve becomes more interesting (as it leaves values near 0). We emphasize that the experimental voltage clamp data is *only* used for model validation, *not* for model creation, considering experimental limitations prevented us from performing the necessary additional experiment (voltage clamp in the presence of an h-current blocker like ZD) to fully isolate the kinetics of the h-current.

The decision to only show these curves in a limited voltage range is a conscious one (although the full model curves are shown later, in Figure 8, to facilitate comparisons). Given the modeling strategy, it would be expected that our h-current model would be most accurate in the voltage regime in which it is most constrained; considering the current clamp traces used, this is roughly the voltage range shown here. Just as we might expect our model neuron to do a poorer job matching the dynamics of the biological neuron outside this range, we similarly expect the h-current model at very hyperpolarized voltages to not be as accurate. This influences the relative inaccuracy of the model’s *τ* value at −90 mV relative to the other data shown in Figure 6, considering that the mathematical model is continuous (i.e. the model *τ* values at very hyperpolarized voltages affect the value at this more moderately hyperpolarized voltage). However, this does not minimize the fact that, just as predicted by the model, experiments show that the approximate kinetics of the h-channel are much slower in human as opposed to rodent neurons. The human experimental data shows maximum *τ* values around 400-500 ms, and the maximum *τ* value in the human h-current model is similar, at approximately 350 ms. However, the Kole et al. (2006) model is different by nearly an order of magnitude, never exceeding 80 ms.

**Figure 8.**
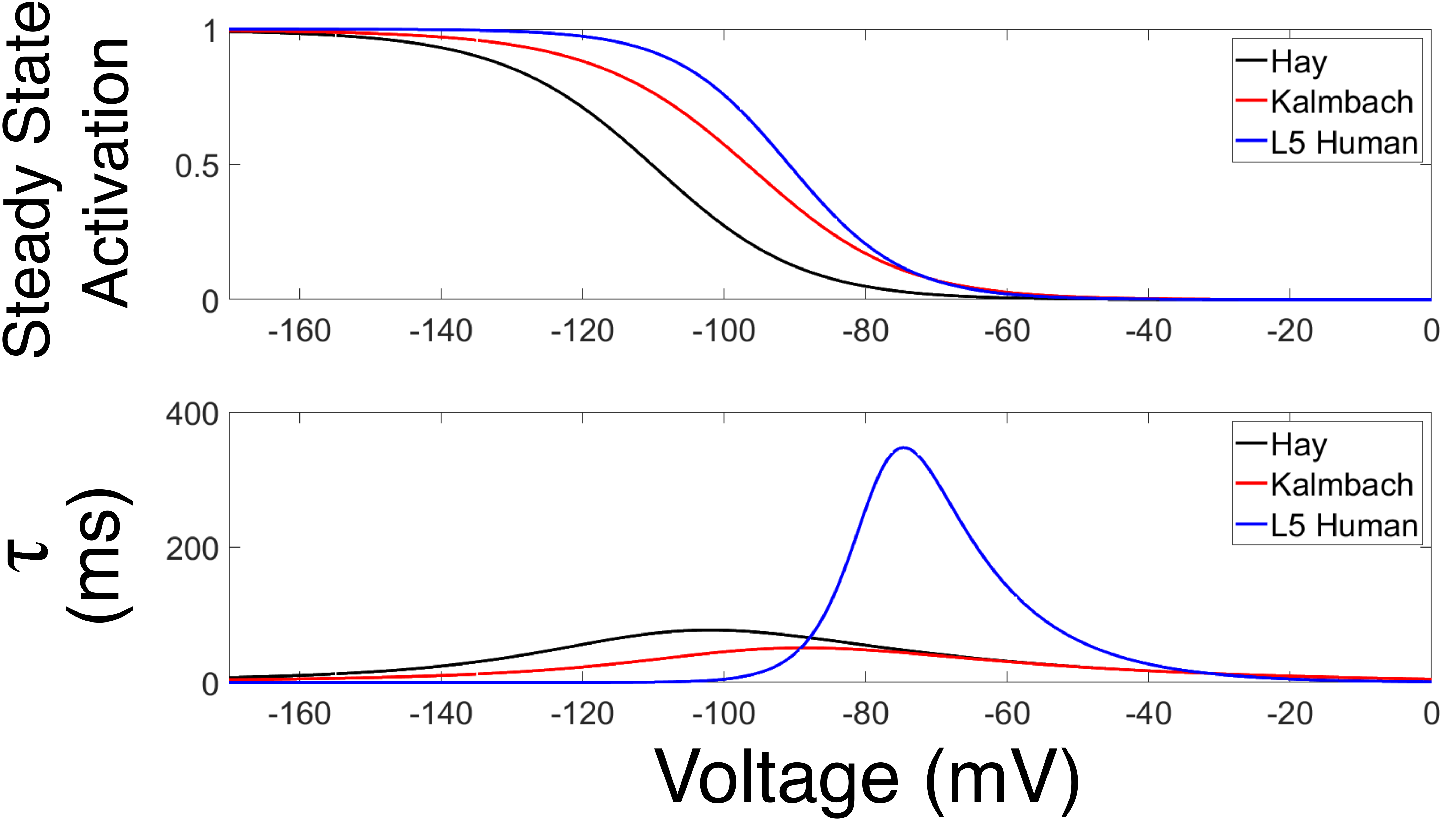
Comparison of h-current models used in three cortical pyramidal neuron models. Plot of steady-state activation curve (top) and *r* (bottom) of the h-current model used by Hay et al. (2011), Kalmbach et al. (2018), and in the model presented in this paper (referred to as “L5 Human”).

While the differences between the human model and the model of Kole et al. (2006) are also apparent in the experimental data alone, no direct comparison of these quantities between the human and rodent settings were made in the corresponding experimental work. Indeed, the computational and modeling work performed here was pivotal in not only highlighting this difference (by comparison between the “human” h-current model and the rodent model of Kole et al. (2006)), but in identifying its functional significance (discussed in great detail in the following). Taken together, these pieces of data validate that our human h-current model is biologically reasonable based on the available experimental results, particularly those from the primary neuron: specifically, the relative magnitude of the *τ* values in our model and the Kole et al. (2006) model lend support to the viability of our model in human L5 neurons.

The fact that the h-current kinetics predicted by our model are biologically reasonable, especially considering their non-trivial divergence from the rodent setting, is pivotal, and justifies our model development approach generally. The fact that we can use mathematical modeling to accurately describe characteristics of h-channels in this setting indicates that the cycling technique described here could be successfully applied to other modeling endeavors where experimental data from a single cell type is similarly limited. In the specific context of this work, these different kinetics and their validation allow for a comparison between rodent and human h-current kinetics. Moreover, considering that h-channels are implicated throughout the literature in determining subthreshold resonance (Kispersky et al., 2012; Zemankovics et al., 2010; Hu et al., 2002; Kalmbach et al., 2018), this model now provides an opportunity to probe the relationship between h-currents and this neural dynamic.

We investigate the model’s capacity for subthreshold resonance by recording the voltage response to the application of a subthreshold ZAP current. We focus on this protocol because data describing the response of human L5 cortical pyramidal cells to this experimental paradigm *in vitro* are presented by Chameh et al. (2019) and so allow comparison. In particular, the human L5 cortical pyramidal cells studied in that work rarely exhibit sub-threshold resonance above 2 Hz. When analogous *in silico* protocols to the experiments presented by Chameh et al. (2019) are performed (described in detail in the Materials and Methods section), our model does not exhibit subthreshold resonance, as shown in Figure 7**A**(in comparison to an example experimental result shown in Figure 7**B**). We note that we will compare these results to those from rodent-derived models in the following section.

This finding provides further validation for our model: despite subthreshold resonance dynamics not being used to directly constrain our model, our model replicates what is seen experimentally under this protocol. This validation extends generally to our modeling approach, as this finding implies that features that were actively “fit” in model generation are essential in driving other, more complex neural dynamics.

Taken together, these validation studies indicate that this model provides a means by which one could explore human-specific dynamics in L5 cortical pyramidal cells. Specifically, an investigation into the relationship between the detailed biophysical model and its various ionic currents (particularly the h-current) and subthreshold behaviours is now well justified. Indeed, the lack of subthreshold resonance observed experimentally by Chameh et al. (2019) was somewhat surprising, as subthreshold resonance (in the 3-5 Hz range) is observed in some super-ficial layer human pyramidal neurons (Kalmbach et al., 2018) and more commonly in rodent L5 pyramidal neurons (Silva et al., 1991; Ulrich, 2002; Dembrow et al., 2010; Schmidt et al., 2016). These experimental results also showed that the “sag” voltage indicating the presence of h-channels is more pronounced in human L5 cells as opposed to deep layer L2/3 (Kalmbach et al., 2018; Chameh et al., 2019). Considering the consensus that the h-current plays some role in driving subthreshold resonance (Hu et al., 2002, 2009; Zemankovics et al., 2010; Kalmbach et al., 2018), these findings might initially seem contradictory. Our model neuron is necessary to probe the potential causal relationship between features of the h-current and subthreshold resonance in detail.

### Inter-species h-channel kinetic differences influence divergent subthreshold resonance characteristics across model neurons

With the model validated, we now compare the behavior of our human L5 cortical pyramidal cell model to two other existing models. The first model is the rodent L5 cortical pyramidal cell model as developed by Hay et al. (2011), which motivated the ion channel types implemented in the human model (see the Materials and Methods). The second model is the human deep L3 pyramidal cell model of Kalmbach et al. (2018), which was built based on human deep L3 morphological and electrophysiological data with the h-channel as the only voltage-gated ion channel type present in the model (see the Materials and Methods for details).

The h-current models used in each of these three models are compared in Figure 8. Moving forward, we will refer to the cell model presented in this paper as the “L5 Human” model to differentiate it from the Hay and Kalmbach models. The differences between our human h-current model compared to the rodent Kole model are that the steady state activation curve is shifted significantly towards more positive voltages, and the kinetics are much slower (indicated by larger values of *τ*), between approximately −90 and −40 mV. In Figure 8, these differences can be seen and compared to the h-current models used by Hay et al. (2011) (the unaltered Kole et al. (2006) model) and by Kalmbach et al. (2018) (a slight adaptation of the model presented by Kole et al. (2006), described in detail in the Materials in Methods).

Given the impetus of this modeling endeavor, we compare the capacity for each of these three models to exhibit subthreshold resonance. In applying an identical ZAP protocol as above for our L5 Human model (see Figure 7), we find that both of these other models, unlike our L5 Human model, exhibit subthreshold resonance at frequencies above 2.8 Hz as shown in Figure 9.

**Figure 9.**
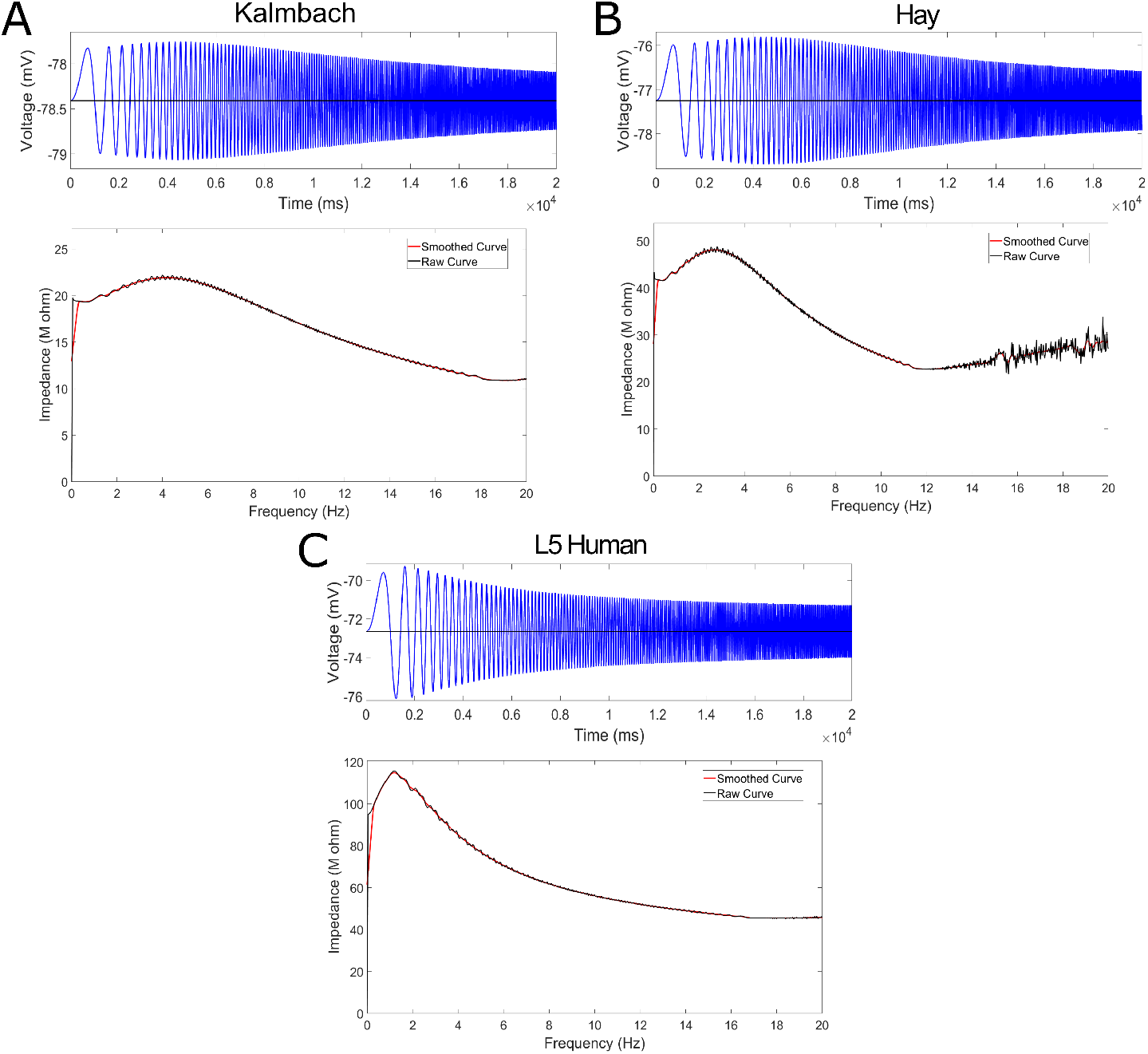
Two pyramidal cell models utilizing h-current kinetics motivated by rodent data each exhibit subthreshold resonance. **(A-C)** Voltage trace (top) and impedance profile (bottom) for the three model pyramidal cells of interest in this study. Previous models from Kalmbach et al. (2018) **(A)** and Hay et al. (2011) **(B)** both exhibit subthreshold resonance, illustrated by a peak in their impedance profiles between 2.8 and 4 Hz. In comparison, the L5 Human model (**(C)**, replicated from previous Figure 7) has a peak barely above 1 Hz.

A quantification of these model comparisons is given in Table 4. Alongside results for the baseline models (illustrated in Figure 9), we also include results for the L5 Human and Hay models with all channels besides the h-channel blocked in order to facilitate a more direct comparison with the Kalmbach model (which has no other active ion channels). This alteration results in a minor change in the resting membrane potential (RMP) of the neuron, as would be expected, but no major change in its peak frequency.

The finding that the Hay model exhibits subthreshold resonance (with a peak in the impedance plot at a frequency near 3 Hz) is as expected considering that subthreshold resonance has been previously observed in rodent L5 cortical pyramidal cells (Silva et al., 1991; Ulrich, 2002; Dembrow et al., 2010; Schmidt et al., 2016). This behavior is also displayed by some of the neurons making up the population studied by Kalmbach et al. (2018), including the neuron motivating their *in silico* model, in which the implemented h-current model was similar to the rodent h-current model presented by Kole et al. (2006) and used by Hay et al. (2011). Indeed, the Kalmbach model also shows subthreshold resonance (with a peak in the impedance plot at 4 Hz). The lack of resonance of our L5 Human model (with the peak in the impedance plot barely above 1 Hz), when contrasted to the subthreshold resonance exhibited by the Hay and Kalmbach models, begs the question of what role the differences in the modeled h-current kinetics might play in dictating this dynamic.

To examine this, we first note that the kinetics of the human h-current model become faster, and in turn more similar to what is seen in the rodent model of Kole et al. (2006) (utilized unaltered by Hay et al. (2011)), at more hyperpolarized voltages (see Figure 8). Thus, if we add a hyperpolarizing DC current to the injected ZAP current to lower the value around which the voltage oscillates, different kinetics for the h-current would also be invoked. It is important to note that, following the discussion related to Figure 6 above, it is highly likely that the h-current’s *τ* value at highly hyperpolarized voltages does not decay to 0 in the biological setting as quickly as it does in our model. For this reason we emphasize that we do not assert that the following experiments make specific predictions about the voltages at which resonance at particular frequencies might occur in the biological setting for human L5 pyramidal neurons. Rather, here we use our existing model as a tool to probe the hypothesis that, given a set morphology, passive properties, and other active channels (as present in our L5 Human model), subthreshold resonance might be dependent on the kinetics of the h-channel being sufficiently fast.

Figure 10 shows the results of such *in silico* experiments for four different values of this hyperpolarizing DC shift. The impedance plots (the bottom figure in each panel) clearly show that, as the mean voltage becomes more hyperpolarized (as can be seen in the top voltage trace plot by a horizontal black line), the curve and the corresponding peak begin shifting rightwards, with an obvious peak beyond 2 Hz appearing in panels **C** and **D**. This resonance is also clearly shown in the corresponding voltage traces. By comparing the different resting voltages in the protocols presented in Figure 10 (and summarized in Table 5) with the voltage-dependent *τ* values in the human h-current model (shown in Figure 8), a correlation is apparent between the tendency to exhibit subthreshold resonance and faster h-current kinetics. Indeed, the resonance is most apparent when the L5 Human model oscillates about voltages where the h-current kinetics are as fast, if not faster, than their rodent counterparts (Figure 10**C-D**).

**Figure 10.**
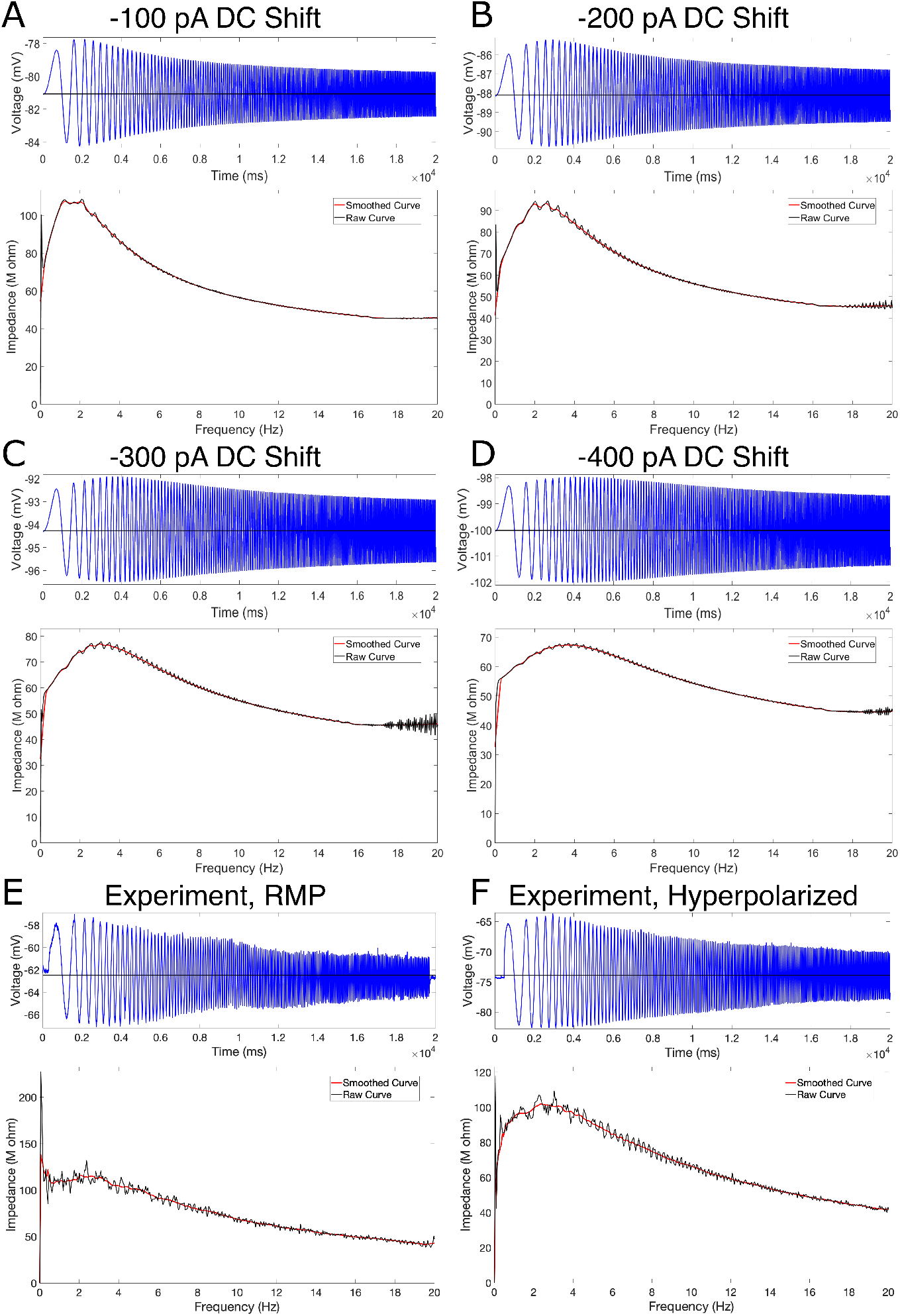
L5 Human model can exhibit subthreshold resonance if held at lower voltages at which the h-current kinetics are faster, implicating these kinetics as playing a crucial role in this dynamic. **(A-D)** Voltage traces (top) and impedance plots (bottom) for ZAP function protocol identical to that shown in Figure 7 **A-B** with the exception of the addition of DC current to hyperpolarize the cell. DC current is −100 pA in panel **(A)**, −200 pA in panel **(B)**, −300 pA in panel **(C)**, and −400 pA in panel **(D)**. Subthreshold resonance reappears clearly as the membrane potential becomes less than −90 mV, where the kinetics of the h-current are as fast or faster than in the Kole et al. (2006) model (see Figure 6). **(E)** Resonance properties of an example human L5 pyramidal neuron experimentally around its resting membrane potential. **(F)** When the same neuron is subjected to a ZAP current when held at a hyperpolarized voltage, more pronounced resonance characteristics are revealed, conforming with *in silico* predictions shown in panels **A-D**.

We note that no such correspondence exists between subthreshold resonance and the steady state activation values of the h-current. At its RMP of −77.25 mV, the Hay model has an h-current steady state activation value below 0.05, while the Kalmbach model has an h-current steady state activation value of approximately 0.15 near it’s RMP of −78.41 mV; both neurons exhibit subthreshold resonance around their resting potentials. Meanwhile, the L5 Human model has an intermediate steady state activation value of approximately 0.075 near its RMP of −72.43 mV. These details illustrate the unlikelihood that the steady state activation values are “confounding” the influence of the h-current’s kinetics, articulated by the *τ* values, on subthreshold resonance. Indeed, if subthreshold resonance was dependent upon having sufficient h-current activity, typified by larger steady-state activation values, we would not expect the neuron whose RMP yields the lowest such value (the Hay neuron) to resonate and a neuron with a higher steady state activation value at RMP (the L5 Human neuron) not to resonate.

While our model h-current’s *τ* value likely decays to 0 at hyperpolarized voltages faster than is observed experimentally, our modeling still makes the prediction that the kinetics of the h-current speed up to some degree below the resting membrane potential. In light of the *in silico* findings described above, this elicits a hypothesis that human L5 pyramidal neurons will exhibit more pronounced subthreshold resonance characteristics when held at hyperpolarized voltages *in vitro*. Preliminary experimental evidence, an example of which is shown in Figure 10**E** and **F**, supports this hypothesis: an example neuron that clearly does not resonate around its RMP shows more pronounced qualities characteristic of subthreshold resonance when the ZAP protocol is applied around a hyperpolarized voltage. Indeed, Figure 10**F** shows a peak in the impedance profile around 3 Hz of similar amplitude to the initial peak near 0 Hz, while Figure 10**E** shows a consistent (albeit noisy) decrease in impedance as the frequency increases with no clear secondary peak away from 0 Hz. We emphasize that this example is meant only as preliminary, anecdotal evidence (indeed, as the resonance properties vary from cell to cell influenced by a variety of properties including morphologies and input resistances, so do their resonance properties around a hyperpolarized voltage); nonetheless, this result provides additional validation of the predictions made utilizing our model. More specifically, this analysis indicates that it is likely the kinetics of the h-current in the human setting *do* speed up when the neuron is hyperpolarized, which *does* result in more pronounced dynamics indicative of subthreshold resonance, even if this effect is likely more gradual than that represented in our model. Note that the voltages shown in the experimental traces in panels **E-F** are not corrected for the liquid junction potential (see Methods).

For comparison purposes, we perform analogous “DC shift” *in silico* experiments on the Hay and Kalmbach models, with the results summarized in Table 5. In each of these hyperpolarized settings both the Hay and Kalmbach models continue to exhibit subthreshold resonance, as would be expected considering such changes do not affect the kinetics of the h-current in these models as significantly as in the L5 Human model.

We emphasize that there are multiple factors at play in determining whether a neuron exhibits subthreshold resonance, not just the activity of the h-current: indeed, the neuron’s morphology, passive properties and other active currents all may play a role (Hu et al., 2002; Kispersky et al., 2012), and the results of Chameh et al. (2019) show that there is heterogeneity in the characteristics of subthreshold resonance even amongst similarly classified human L5 pyramidal cells. However, we note that using our *in silico* model we are able to more directly address the contribution of the h-current in the neurons’ responses to these protocols. In particular, when comparing the L5 Human and Hay models (given that the Kalmbach model only contains the h-current), we find that the h-current is the dominant inward ionic current when the ZAP current is delivered alongside a hyperpolarizing DC current, which is not surprising given the known voltage dependence of the various ionic currents modeled here (see the full equations dictating the various ionic currents’ voltage dependencies in Hay et al. (2011)). Even at the resting membrane potential, as shown in Table 4, blocking other active currents does not affect the L5 Human neuron’s capacity for resonance, allowing us to reasonably assume that the activity of currents besides the h-current are not playing a major role in dictating this neuron’s lack of subthreshold resonance. We note that this blockade, and the ability to identify the contributions of specific ionic currents to the voltage response to a ZAP current, takes advantage of our computational models and analyses.

In our endeavor to support the hypothesis that a relationship exists between the kinetics of the h-current and a neuron’s capacity for subthreshold resonance, the above analysis provides support in one logical “direction”: by “speeding up” the kinetics of the h-current in the setting of our L5 Human model, resonance is observed where it previously was not. If we can provide support in the other “direction”, namely by showing that “slowing down” the kinetics of the h-current can eliminate resonance where it once was present (i.e. the Hay or Kalmbach models), we will have more complete logical support of our hypothesis. We perform such an investigation via an examination of “hybrid” neural models in which rodent h-current models (that of Hay and Kalmbach) are replaced with the human h-current model; in doing so, the *only* change in a “hybrid” model from its original state is in the kinetics of the h-current. We note that this investigation is likely only possible *in silico*. This choice not only achieves the desired logical goal, but also allows for potentially broader conclusions to be drawn regarding human and rodent differences.

Before beginning this investigation, it is important to note that such a switch between human and rodent h-current models would affect other aspects of the cellular model (including, for example, the resting membrane potential, as well as the potential activity of other ion channels) that might affect its behavior. Moreover, the differing morphology and passive properties that make up the “backbones” of these models also differ significantly, and these properties also play a role in dictating a neuron’s frequency preference (Hutcheon and Yarom, 2000; Rotstein and Nadim, 2014). It is for these reasons that we emphasize that, in performing such a “switch”, we create new “hybrid” models that must be approached cautiously. However, a very specific focus on the subthreshold dynamics of these “hybrids” makes their use as presented here reasonable. There are two primary rationales for this assertion: first, a focus on subthreshold dynamics significantly minimizes the role that other ionic currents (whose features vary between “model backbones”) will play in the dynamics; and second, by only switching the h-current models (i.e. the kinetics of the h-current), and not the distribution nor conductance of the h-channel, the focus can be mainly on how the different kinetics might play a role (i.e., differences shown in Figure 8).

The results obtained are summarized in Table 6. Most critically we observe that, when the Hay and Kalmbach models have their respective h-current models replaced with the human h-current model, these “hybrids” have either a clear lack of resonance (Hay-L5 Human Hybrid) or a peak frequency now below the typical “theta” range (Kalmbach-L5 Human Hybrid). As the RMPs of these “hybrids” are within the range of voltages for which the human h-current displays significantly slower kinetics than the rodent models, these results are support for the second “direction” in our argument: namely, by “slowing down” the h-current kinetics in the hybrid model as compared to the baseline model, we severely diminish (if not entirely eliminate) the previously observed subthreshold resonance. Doing so in this fashion also further emphasizes the importance of the differences in the human and rodent h-current models in dictating neural dynamics.

For completeness, we perform analogous experiments with a DC shift on these hybrids as was done on the L5 Human model. As expected, in the “hybrids” in which a rodent h-current model is replaced by the L5 Human h-current model, a hyperpolarizing DC shift can serve to reestablish subthreshold resonance, just as in the baseline L5 Human pyramidal cell model. Indeed, with −400 pA DC shifts, both the “Hay-L5 Human” and the “Kalmbach-L5 Human” models show a preferred frequency greater than 2 Hz, and the hyperpolarized resting voltages under these protocols are in a range at which the kinetics of the human h-current approach the kinetics of the rodent h-current models.

We note here that these results are further evidence that the steady state activation values of the h-current do not play a major role in dictating subthreshold resonance, while the *τ* values do. Indeed, in both hybrid models the RMP is at or below that seen in the default L5 Human model, yielding a larger steady state activation value for the h-current and thus more h-current activity; however, this does not lead to more pronounced resonant activity. Thus, we remain confident that the feature of the h-current kinetics of primary importance in a neuron’s capacity for subthreshold resonance is the speed of its activity (i.e. the *τ* value) rather than the amount of steady state activity (i.e. the steady state activation value).

Taken together, these results provide strong support for the argument that the differing h-channel kinetics in L5 between humans and rodents play a paramount role in dictating the neural dynamic of subthreshold resonance. This support is bolstered by the dual directions of our causal argument: we can “rescue” resonance by “speeding up” h-channel kinetics, and we can “eliminate” resonance by “slowing down” h-channel kinetics. The additional fact that eliminating resonance can be achieved by “slowing down” h-channel kinetics by imposing our human h-current model on a rodent L5 pyramidal cell model, thus creating a “hybrid” model, further emphasizes the functional importance of the inter-species differences identified both experimentally and computationally.

## Discussion

In this work, we present a biophysically detailed, multi-compartment, full spiking model of a human L5 cortical pyramidal cell that is constrained primarily from morphological and electrophysiological data from the same neuron. The model leads to a mathematical characterization of h-currents that is specific to human cortical cells and is validated against experimental data from the primary neuron that was not used in model development. Our model additionally mimics subthreshold (a lack of resonance) characteristics observed experimentally in a separate population of human L5 cortical pyramidal cells (Chameh et al., 2019). Moreover, the model replicates the capacity for PIR spiking in these cells that is thought to be driven by h-channels (Chameh et al., 2019), and encapsulates other general spiking characteristics of these neurons. This indicates that our fitting procedure was able to capture a crucial “essence” of these cells’ complex dynamics, even given the challenges posed in this modeling endeavor by the current limitations of human electrophysiological data in comparison to the rodent setting.

This unique computational model allowed us to perform a detailed *in silico* investigation into the relationship between subthreshold resonance and the h-current; many of the *in silico* experiments performed in this manuscript would be infeasible or intractable *in vitro*. This exploration provided convincing support of a strong relationship between the time constant of the h-current’s activity and the capacity for subthreshold resonance. Such resonance can be “rescued” in cells in which it is absent by “speeding up” h-current kinetics, and “eliminated” in cells in which it is present by “slowing down” h-current kinetics. This relationship indicates that there are key functional consequences to inter-species biophysical, cellular differences like those identified in this research.

### Modelling goals, strategies and comparisons

All computational models are an idealization and abstraction of the physical entity of interest. Given the inherent limitations on such modeling endeavors, the choices of where the necessary approximations are implemented must be made with an overall research question in mind. Such choices should ensure that it is reasonable to use the model to make inquiries into the particular question of interest, which may come at the cost of the model’s applicability in other contexts. Indeed, it is highly unlikely given contemporary tools that an entirely “realistic” neuron model, encapsulating all known properties and dynamics of a biological cell, can ever be obtained; instead, computational neuroscientists must limit the scope of their inquiries and conclusions to the context in which the model was constrained, and is thus the most “realistic” (Almog and Korngreen, 2016).

Here, we used a unique data set, namely morphology and a suite of current clamp recordings (in the presence of TTX) obtained from the same human neuron, in the development of a biophysically detailed human neuron model that could help us understand the distinctness of human brain dynamics. By primarily constraining our model with these data, we minimized the likelihood that cell-to-cell variability could compromise the model (Marder and Goaillard, 2006; Golowasch et al., 2002). However, naively “fitting” our model to just these current clamp recordings omitted a crucial component of the neuron’s function: its spiking characteristics. Given that all recordings from our primary neuron were obtained in the presence of TTX, we could not infer any such characteristics from this primary neuron. This led to the implementation of the informed “cycling” fitting technique schematized in Figure 2**A**. In this fashion, we maintained the benefits of the primary constraining data coming from the same neuron, while also ensuring the neuron retained general spiking characteristics of similarly classified neurons, with an emphasis on PIR spiking.

This strategy was motivated in part by the discussion presented by Roth and Bahl (2009) and cyclical “re-fitting” strategies (Sekulić et al., 2015, 2019). The particular approach we developed here to use with human experimental cellular data could be useful for future modeling endeavors with limited experimental data from a single neuron. We emphasize that using data from the same cell mitigates the potential impact that averaging values, such as passive properties, over multiple cells might have on our model. Indeed, it is well established that the morphology of the neuron plays an important role in dictating its passive properties (Mohan et al., 2015; Eyal et al., 2016; Beaulieu-Laroche et al., 2018; Gouwens et al., 2018); as such, imposing passive properties obtained from multiple neurons onto a single morphology is fraught with the potential for error. This would extend to the h-current model and other ionic current models imposed on the passive “backbone” of the reconstructed neuron; specifically, properties of the h-current are particularly vulnerable to such errors considering the h-channel’s non-uniform distribution along the dendrites (Kole et al., 2006; Ramaswamy and Markram, 2015; Beaulieu-Laroche et al., 2018).

It is worth highlighting that our modeling approach was designed specifically to make best use of the data set obtained from our primary neuron and presented in this work, along with a consideration of the goals of this research. Given that the amount of human data is incredibly scarce as compared to rodent, it did not seem appropriate at this time to pursue a “model database” exploration to examine aspects such as compensatory biophysical mechanisms (Marder and Taylor, 2011). However, the modeling approach articulated here may provide an undergird for future attempts to create such model databases of human neurons as human electrophysiological recordings continue to expand. Additionally, given our unique data set we chose to constrain this model by directly “fitting” the voltage responses of the model neuron to current clamp protocols to what was observed experimentally, with the choice of these protocols informed by the features of primary interest in our study. In contrast, many similarly detailed rodent models (including those discussed below (Hay et al., 2011; Dong, 2008; Jones et al., 2009; Sunkin et al., 2012)), use a “feature fitting” technique that fits quantifications of more complicated neural behaviors in the model to that seen experimentally. As both of these techniques have been shown to yield useful models and biological insights, the choice of modeling technique is driven by the available data (i.e. the differences in data availability in the rodent and human setting) and the goals of the study. Indeed, it was only through directly fitting our neuron to current clamp recordings that we were able to articulate our novel model of the dynamics of the human h-current.

We compared our human L5 cortical pyramidal cell model with two existing models: the multi-compartment, rodent L5 cortical pyramidal cell of Hay et al. (2011), and a multi-compartment model of a human cortical deep L3 pyramidal cell with only passive properties and the h-current presented by Kalmbach et al. (2018). Each of these models provides a useful point of comparison, the Hay et al. (2011) model because it is of an analogous rodent neuron with similar computational detail, and the Kalmbach et al. (2018) model because it is constrained by human data. We note that we chose to compare our model to the human model presented by Kalmbach et al. (2018), rather than another human L2/3 model presented by Eyal et al. (2016), considering that the work of Kalmbach et al. (2018) specifically focuses on the h-current and subthreshold resonance, and thus provides a direct point of comparison given the goals of this work.

The Hay et al. (2011) model informed the choice of ion channels implemented in our model given that it was also of a L5 pyramidal cell. During model generation we found that a best “fit” to our human experimental data led to significant changes in a variety of maximum conductances (see Table 1) as well as the h-current kinetics. We note that there exist a variety of other L5 rodent cortical pyramidal cell models (Keren et al., 2009; Almog and Korngreen, 2014; Farinella et al., 2014; Larkum et al., 2009) that are focused on features, often concerning spiking behavior, observed in rodent neurons. Thus, while these models may be better suited for *in silico* investigations of these neural dynamics generally speaking, our developed model, based on human data, is more appropriate to use for an investigation of human cortical behaviors.

Here it is worth noting the potentially surprising result that our maximum conductance value for the h-current is lower than that used in the model of Hay et al. (2011) (see Table 1), despite experimental evidence of larger sag currents in human neurons (Kalmbach et al., 2018; Chameh et al., 2019). However, our model still matches the large sag exhibited by human L5 neurons. This is due to the novel human h-current model, which includes larger steady-state activation values at these hyperpolarized voltages than the Kole et al. (2006) rodent model implemented by Hay et al. (2011). Indeed, this maximum conductance value alone does not dictate the “amount” of h-current active in the model neuron.

The comparison between our model and other human neuron models is less clear than the conspicuous rodent versus human difference, although the number of these models is limited by access to human tissue. Beaulieu-Laroche et al. (2018) present a human L5 cortical pyramidal cell model, but unlike our current work, its morphology was not directly based on a human pyramidal cell. Rather, a modified rat pyramidal neuron morphology was “stretched” to allow comparison to the rodent model of Hay et al. (2011). We also note that the model presented by Beaulieu-Laroche et al. (2018) uses the rodent h-current model of Kole et al. (2006), so that its h-current driven dynamics would likely be quite similar to the models studied in-depth here (that of Kalmbach et al. (2018) and Hay et al. (2011)) that do have a detailed biophysical morphology. We further note that while the Allen Institute is one of few laboratories currently using human data to generate computational neuron models with the level of morphological detail presented here, the human models that are presently a part of the Allen Brain Atlas (Dong, 2008; Jones et al., 2009; Sunkin et al., 2012) have their voltage-gated ion channels confined to somatic regions. In contrast, the model presented here has voltage-gated ion channels distributed throughout the dendrites, following that of rodent ion channel distributions. The recent model presented by Kalmbach et al. (2018) moves toward the expression of ion channels in dendritic regions, as h-channels are included throughout the dendrites. However, as mentioned above, the only voltage-gated channels included in that model are h-channels.

A more recent model of a human L2/3 neuron by Gidon et al. (2020), which takes advantage of dendritic recordings in human neurons like Beaulieu-Laroche et al. (2018), utilizes a detailed morphology of a human L2/3 pyramidal cell and instantiates a “phenomenological” implementation of calcium-mediated dendritic action potentials (dCaAPs). This is used to show that these neurons can potentially solve complex computations that are thought to require multi-layer neural networks. Thus, the model of Gidon et al. (2020) argues that the specific characteristics of human cells matter in driving potentially functionally significant dynamics. This can be seen as parallel to the results of this work, in which we find that the emergent dynamic of subthreshold resonance is dependent on the biophysical characteristics of human h-currents. It is worth emphasizing that the modeling strategies underlying these two works are complementary: these models occupy distinct yet non-competitive spaces in the computational literature, both moving the field of human neuronal modeling forward and answering separate research questions.

### H-channels, resonance and function

H-channels are tetramers that can be either homomeric (consisting entirely of the same subunit type) or heteromeric (consisting of different subunit types) (Biel et al., 2009; Shah, 2018). Interestingly, one of the primary differentiating factors between the four subunits are their time constants of activation, with HCN1 subunits being the fastest, HCN4 being the slowest, and HCN2-3 lying in between (Shah, 2018). Viewed in the context of our study, the slower kinetics that we observe both computationally and experimentally in human L5 pyramidal cells in comparison to their rodent counterparts suggests that human L5 pyramidal cells might have enriched non-HCN1 subunits amongst their h-channels. Indeed, human neurons in general, and L5 pyramidal cells specifically, have an enrichment of HCN2 channels as revealed via mRNA expression (Kalmbach et al., 2018). HCN2 subunits have slower activation kinetics and a more negative half-activation voltage than HCN1 subunits (Biel et al., 2009), with research showing that heteromeric h-channels consisting of a mix of HCN1 and HCN2 subunits display slower kinetics than those seen in HCN1 homomeric h-channels (Chen et al., 2001). Taken together, these results suggest that the differences in h-channel kinetics may be driven by different HCN subunit expression between rodent and human L5 pyramidal cells. A detailed comparison between subunit expression in rodents and humans remains wanting given the clear predictions of this study. Additionally, since channel kinetics can be altered by post-translation modification, proteomics may be helpful in investigating post-translation modification of HCN subunits in human neurons.

The pacemaking and resonant contributions of h-channels have led to them being a focus of many studies (Biel et al., 2009). In particular, the role played by h-channels in dictating subthreshold resonance properties has been examined in excitatory cells (Hu et al., 2002, 2009; Kalmbach et al., 2018; Zemankovics et al., 2010; Silva et al., 1991; Ulrich, 2002; Dembrow et al., 2010; Schmidt et al., 2016), as well as inhibitory cells (Kispersky et al., 2012; Zemankovics et al., 2010; Sun et al., 2014; Stark et al., 2013) both in hippocampus and cortex, and the frequency of this subthreshold resonance has been found to be in the theta frequency range (3-12 Hz). This makes the difference between human and rodent h-channel kinetics and subthreshold resonance of further interest, since differences between human and rodent theta rhythm frequencies have been noted (Jacobs, 2014), with traveling theta waves in human hippocampus and neocortex shown to be important in human cognition (Zhang and Jacobs, 2015; Zhang et al., 2018). However, the relationship between subthreshold and suprathreshold resonant and oscillatory dynamics has yet to be fully articulated: for example, a given subthreshold resonant frequency does not necessarily lead to a similar spiking resonant frequency (Rotstein and Nadim, 2014; Rotstein, 2017). The dendritic filtering capacities of neurons (e.g., see Vaidya and Johnston (2013)) adds another layer to this relationship. Theoretical and computational studies bring forth the importance of understanding the complexity of the interacting dynamics from different ion channel types and the passive properties in pursuit of better understanding this relationship (Hutcheon and Yarom, 2000; Rotstein and Nadim, 2014; Rotstein, 2017; Sekulić and Skinner, 2017).

### Limitations and future work

The Hay et al. (2011) model informed the choice of ion channels implemented in our human L5 model given that it was also of a L5 pyramidal cell. In this vein, we note that the non-uniform distribution of h-channels implemented in our model is driven from rodent findings (Kole et al., 2006). While there is some experimental evidence that h-channels are similarly distributed in human neurons (Beaulieu-Laroche et al., 2018), it is likely that there are some differences in these distributions given the distinct morphologies of similarly classified rodent and human pyramidal neurons. Thus, we followed the distribution of rodent h-channels in this model as a necessary strategy given the absence of similarly detailed human data. This is an aspect of the model that may be improved upon as such data becomes available.

As with any modeling endeavor, there are limitations on the contexts in which the model can be appropriately used. Although our model was not constrained by spiking properties such as backpropagating action potentials or calcium spikes like the Hay et al. (2011) rodent model, this choice was motivated by the overall focus in this study on h-channel driven dynamics. The spiking characteristics constraining model development were limited to repetitive spiking frequencies and the capacity for PIR spiking observed in data from other human L5 pyramidal cells (Chameh et al., 2019). Thus, any investigation of suprathreshold characteristics of this model must be done with the important caveat that such constraining data did not come from the primary neuron used in model creation. Furthermore, other features of cortical pyramidal cells that might influence the dynamics of human L5 pyramidal cells, such as the spike shape (Molnár et al., 2008), calcium spiking (Hay et al., 2011), backpropagating action potentials (Hay et al., 2011; Larkum et al., 1999) and synaptic responses (Molnár et al., 2008; Eyal et al., 2018) that were not used in model creation should be considered carefully in future studies. However, considering that our model captured inter-species differences and showed their influence on subthreshold resonance, we believe that the model presented here is more suitable than rodent models for an investigation of distinctly human cortical neuron dynamics.

Before using the presented model in novel settings, it would be helpful to perform additional “confirmations” to gauge whether the human-specific properties of interest are displayed by the model. For example, in characterizing a large population of human L5 cortical pyramidal neurons, Chameh et al. (2019) investigated both subthreshold and suprathreshold dynamics. The frequency-dependent gain measure developed by Higgs and Spain (2009) encapsulates a cell’s phase preference for spiking in response to an oscillatory input as a function of frequency. While such suprathreshold behaviors were not a focus of the present modeling endeavor, given that we now have a full spiking model, we can consider our model in light of this experimental data and gain further insight into human brain function. We have applied an analogous *in silico* protocol to our model neuron and are encouraged that key features of the frequency-dependent gain are indeed captured by this model, including the presence of a “peak” in the delta range that is dependent upon h-channel activity and a “trough” in the 5-10 Hz range (these results are not shown here as they are the focus of future work).

Further, in contexts where the model presented here is not immediately appropriate, “adjustments” based on other experimental data can be made to answer different research questions, just as was done by Shai et al. (2015) in their adjustments to the Hay et al. (2011) model. Indeed, such research is a fertile ground for future work utilizing this model: one potential avenue is better encapsulating the medium afterhyperpolarization (mAHP) implicated in determining a neuron’s suprathreshold frequency preference (Higgs and Spain, 2009) in order to make the model appropriate for an *in silico* investigation into the different influences the h-current and the mAHP play on these spiking features. Another is an investigation of the potential for back-propagating action potentials in this distinctly human model, a feature studied in detail in the rodent model of Hay et al. (2011).

Finally, of interest is the relationship between h-channels, PIR, and epilepsy. Recent experimental (Chang et al., 2018) and computational (Rich et al., 2020) support has been presented for a novel hypothesis of seizure initiation, termed the “GABAergic initiation hypothesis”, which posits that excessive inhibitory signalling may trigger seizure via a cascade of events including PIR spiking in pyramidal cells. The experimental work of Chameh et al. (2019) reveals that human cortical L5 pyramidal cells can exhibit PIR spiking under current-clamp conditions, likely driven by the dynamics of the h-channel. By including this important dynamic as part of the “biological tether” of the research presented here, this model may prove useful in an investigation of whether similar PIR spiking is viable *in vivo*.

In conclusion, given the preciousness of human data and the need to acquire as much data from the same cell to appropriately create specific cell type models, we have developed a strategy by which distinctly human neuron models can be created. We have built a human L5 cortical pyramidal cell that has been validated and used it to provide an explanation for human-specific subthreshold resonance dynamics.

## Acknowledgments

We thank Happy Inibhunu for applying new data visualization techniques to this model that helped to shape the presentation of these results. We thank Costas Anastassiou, Anatoly Buchin and Tom Chartrand of the Allen Institute for their open communication and assistance facilitating the use of their model in this work. We thank Etay Hay for providing critical feedback on this paper during the early writing stages. We thank Clayton S. Bingham for his valuable feedback on our model and for fruitful discussions regarding its potential applications.

## Corresponding Author Address

Krembil Research Institute, Krembil Discovery Tower, Toronto Western Hospital, 60 Leonard Avenue, 4KDT489, Toronto, ON, Canada, M5T 0S8

